# Encoding of task regularities links grid-like signals to human timing behavior

**DOI:** 10.1101/2024.01.23.576694

**Authors:** Ignacio Polti, Matthias Nau, Raphael Kaplan, Virginie van Wassenhove, Christian F. Doeller

## Abstract

Grid cells have been proposed to encode task regularities that allow predicting future states. Entorhinal grid-like signals might therefore mirror behavioral biases associated with relying on task regularities, like regression-to-the-mean biases in time estimation. Here, we tested this proposal using functional magnetic resonance imaging and a rapid timing task in humans. Indeed, trial-wise entorhinal activity reflected task accuracy and the degree to which interval estimates regressed towards the mean of all tested intervals. Grid-like signals were observed exclusively for the interval closest to the mean, which was explained by differences in temporal grid stability across intervals. Finally, both behavioral and entorhinal results were explained by a Bayesian observer model that assumes the integration of current-trial sensory evidence with prior expectations. Together, we find that entorhinal activity and grid-like signals reflect behavioral performance in a timing task, supporting the proposed role of grid cells in encoding task structure for predictive processing.

## Introduction

The ability to recognize and utilize statistical regularities governed by the co-occurrence of stimuli, actions, and events is crucial for any successful interaction with the environment (Vetter, Wolpert, 2000; Clark, 2013; Friston, Buzsáki, 2016). Learning about such regularities allows us to predict future states of the world, for instance to track moving objects during occlusion, which is an essential ability underlying flexible and robust behavior broadly (Fiser et al., 2010; Heald et al., 2023; Schapiro, Turk-Browne, 2015; Schapiro et al., 2016). When catching an approaching ball, for example, we anticipate the moment it will reach us not solely based on our estimates of its current speed and distance, but also based on our knowledge of how previous balls have behaved in similar situations. Through experience, we have learned about the probability associated with certain speeds and arrival times, which now guides when and how we act. How does the brain encode such statistical regularities and how do they afford predictive inference in the service of behavior?

A prominent metaphor of how the brain encodes statistical regularities can be found in the concept of cognitive maps, referring to relational map-like representations of places and events that support mnemonic and predictive processes (O’Keefe, Nadel, 1978; Moser et al., 2014; Stachenfeld et al., 2017; Whittington et al., 2020; Eichenbaum et al., 1999). A large body of neuroscientific literature on cognitive maps and predictions pointed to the hippocampal formation as a key neural component involved in both, fueling efforts to unify theories of its contribution across a range of tasks (Whittington et al., 2020; Stachenfeld et al., 2017; Ambrogioni, Ólafsdóttir, 2023; Behrens et al., 2018; Bellmund et al., 2018). An emerging view is that the hippocampal formation plays an important role in encoding regularities that afford generalization across tasks or contexts (e.g., different environments; Whittington et al. (2020); Bousquet et al. (1998); Fuhs, Touretzky (2007); Penny et al. (2013); Friston, Buzsáki (2016); Pezzulo et al. (2017)), thus greatly accelerating learning and reducing behavioral errors in novel or noisy situations (Lisman, Redish, 2009; Stachenfeld et al., 2017). In particular, the hippocampus has been suggested to support the encoding of task regularities in real time as a task is performed, as its activity reflects feedback and behavioral improvements even in fast-paced timing tasks (Polti et al., 2022).

Like the hippocampus, the entorhinal cortex is widely considered critical for cognitive mapping; not only is it a major anatomical gateway for hippocampal-cortical interactions (Witter, Amaral, 2004), but it also harbors grid cells (Hafting et al., 2005; Moser et al., 2014) thought to represent relationships between places and events in the world (e.g., different spatial positions during navigation, but also non-spatial features (Constantinescu et al., 2016; Aronov et al., 2017; Behrens et al., 2018; Bellmund et al., 2018; Bao et al., 2019; Theves et al., 2019, 2020; Viganò, Piazza, 2020; Park et al., 2021). This grid-like “coordinate system” may be predictive in nature, meaning that it likely anticipates future states of the agent (e.g., future positions during navigation, Stachenfeld et al. (2017)) and affords efficient vector computations for spatial planning (Banino et al., 2018; Bush et al., 2015; Bicanski, Burgess, 2020). At the population level, grid-cell activity is believed to exhibit a hexadirectional (i.e., six-fold rotationally symmetric) modulation as a function of virtual running or gaze direction (Killian et al., 2012), which can be observed in the human entorhinal cortex using functional magnetic resonance imaging (fMRI) (Doeller et al., 2010; Julian et al., 2018; Nau et al., 2018b; Graichen et al., 2025). Importantly, while grid-like signals have been observed in a range of tasks and species (Kunz et al., 2019), most studies left open whether or not they were indeed relevant for task performance.

Here, we set out to understand the contributions of the entorhinal cortex and grid-like signals in particular to the encoding of task regularities that afford predictive coordination of actions relative to sensory events. In addition, we investigated the relationship between grid-like signals and task performance. For this purpose, we monitored human brain activity using fMRI while participants engaged in a visual tracking task previously shown to engage grid-like signals in the human entorhinal cortex (Nau et al., 2018b) as well as a time-to-contact (TTC) estimation task shown to engage the adjacent hippocampus in a rapid and behavior-dependent manner (Polti et al., 2022). We specifically tested whether and how entorhinal activity reflects behavioral biases previously linked to the encoding of timing-task regularities (Jazayeri, Shadlen, 2010), focusing on the posteromedial portion of the entorhinal cortex (pmEC); the presumed human homologue of the rodent medial entorhinal cortex that harbours grid cells (Navarro Schröder et al., 2015; Maass et al., 2015; Syversen et al., 2021).

## Results

In the following, we will present our experiment and results in 5 steps. First, we introduce our task design and empirical measures in detail. Second, we show that task performance in each trial depended on the intervals tested in previous trials, with behavioral responses showing a systematic bias towards the average interval. This regression-to-the-mean bias suggests that participants relied on prior knowledge accumulating across trials. Third, we report that trial-wise pmEC activity mirrored this behavioral bias in real time, similar to previous reports for the hippocampus (Polti et al., 2022). Fourth, we show that human pmEC grid-like signals co-varied with the tested intervals across trials. This effect was explained by the temporal stability of the grid-like signal (i.e. replicability across data partitions), and its amplitude was correlated with behavioral performance in all participants. Finally, by showing that a Bayesian observer model provides a parsimonious account of the data, we illustrate a potential computational explanation of our results, namely that entorhinal grid-like signals may reflect the mismatch between prior knowledge and sensory evidence obtained in each trial. Collectively, these results suggest that non-spatial task factors (such as tested intervals) shape spatial representations in the entorhinal cortex in service of timing behavior. Moreover, we provide evidence that entorhinal activity, and grid-like signals in particular, reflect the rapid encoding of task regularities in service of immediate task performance.

### Time-to-contact (TTC) estimation task

Our task consisted of two components; a visual tracking task that engages grid-like signals in the human EC (Nau et al., 2018b), as well as a predictive timing task that engages other regions including the adjacent hippocampus (Polti et al., 2022). Over the course of 768 trials, participants tracked a moving fixation target with their eyes until it was occluded, which then prompted them to predict when the target would hit a visual boundary. In each trial, the fixation target moved 10 degrees of visual angle (dva) into one of 24 directions (Fig. 1A, “Gaze trajectory”) at one of 4 possible speeds, yielding 4 different intervals to be estimated (*TTC*_*t*_): 0.55, 0.65, 0.86, and 1.2 s (See *Methods*). Speed and direction were held constant within each trial. After the target stopped moving, participants estimated the time when it would have hit the visual boundary 5 dva apart, which they indicated via button click (Fig. 1A, “TTC estimation”). Participants then received feedback reflecting the accuracy of their estimated TTC relative to the ground-truth TTC. The next trial then started after a jittered inter-trial interval (ITI). See the methods section for details.

**Figure 1:**
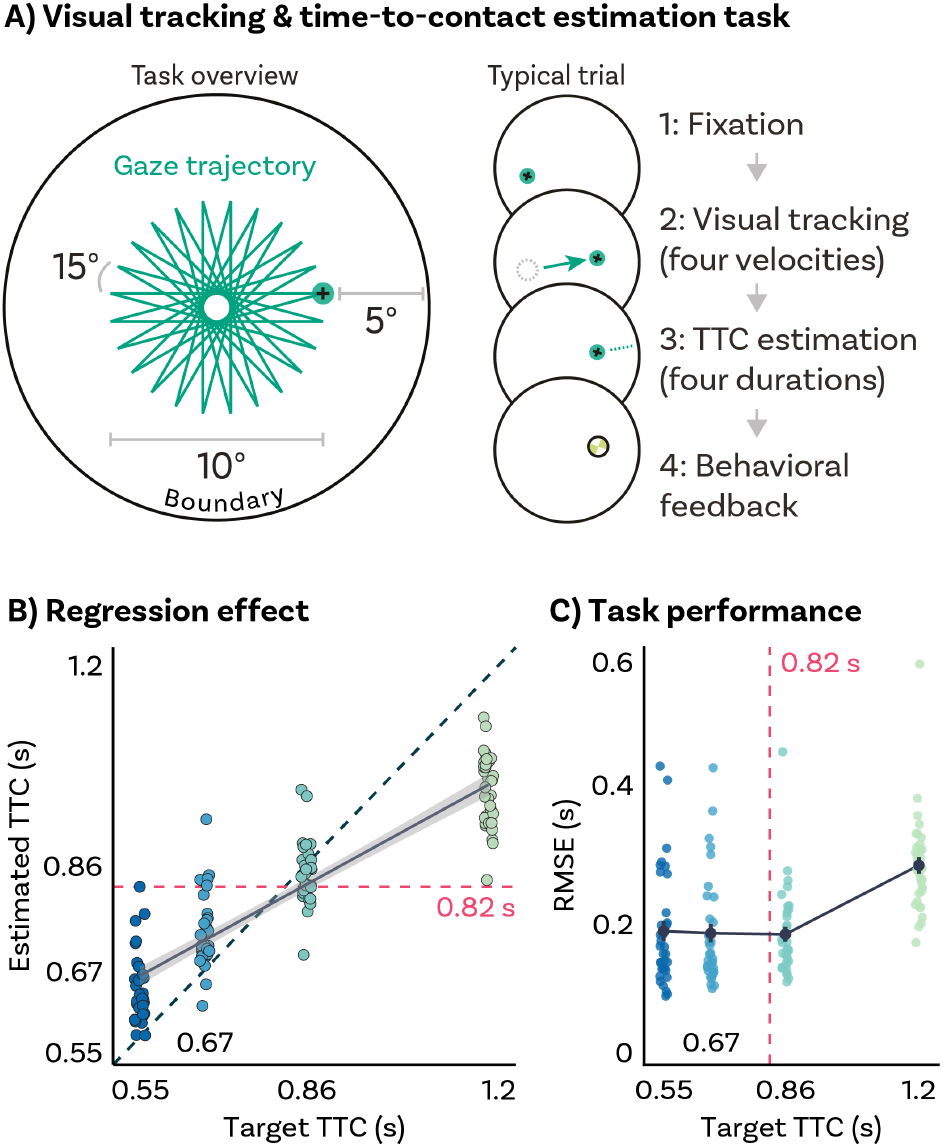
A) Visual tracking and Time-To-Contact (TTC) estimation task. In each trial during fMRI scanning, participants fixated on a target (1), which started moving at one of 4 possible speeds and in one of 24 possible directions for 10 dva (2). After the target stopped moving, participants kept fixating and estimated when the fixation target would have hit a boundary 5 dva apart (3). After pressing a button at the estimated TTC, participants received feedback (4) based on their performance. B) Regression effect. Participants responses regressed towards the mean of the sampled TTCs (0.82 s, horizontal dashed line), away from the identity line (diagonal dashed line). Regression line (black) and standard error (gray shade). C) Task performance. Root-mean-square error (RMSE) differences across TTCs show a quadratic pattern centered on the mean of the sampled TTCs (0.82 s, vertical dashed line) have a lower RMSE. We plot the mean and SEM (black dot and line). BC) Single-participant data plotted as dots. Target TTCs are color coded.

### Behavioral results

To examine whether participants performed the task well, we first compared their estimated TTC (*TTC*_*e*_) to the target TTC (*TTC*_*t*_, ground truth) using a mixed-effects model (MEM). Indeed, we found that the estimated and target TTCs were tightly correlated (Fig. 1B; *F*(1, 32.99) = 476.83, *p* = 2.2*x*10^−16^, *ϵ*^2^ = 0.93, *CI*: [0.90, 1]). However, TTC estimates were further systematically biased towards the mean of the tested intervals (0.82 s, Fig. 1B, horizontal dashed line) in line with previous reports using timing tasks (Miyazaki et al., 2005; Jazayeri, Shadlen, 2010; Acerbi et al., 2012; Cicchini et al., 2012; Chang, Jazayeri, 2018). We quantified this regression-to-the-mean bias by extracting the MEM *TTC*_*t*_ group-level coefficient, which resulted in a slope value of 0.53 (Fig. 1B, MEM fit, diagonal solid line). For comparison, perfect task performance would lead to a slope of 1, whereas total regression to the mean would result in a slope of 0.

To quantify task performance in more detail, we then calculated the variability and bias in TTC estimation for each participant and tested interval. Variability describes how similar the estimates were across trials, whereas bias describes how similar they were to the ground-truth *TTC*_*t*_ (see methods for details). We found that variability increased as a function of *TTC*_*t*_, and that bias inscreased with the distance of each *TTC*_*t*_ from the mean of the sampled TTCs (Fig. S1A). Together, these two measures combine into the root-mean-square error (RMSE), which we computed as our summary performance measure. We found that the RMSE showed a quadratic relationship to the *TTC*_*t*_, centered on the mean of the sampled *TTC*_*t*_ (0.82 s; Fig. 1C; MEM, *F*(2, 35.36) = 58.37, *p* = 6.29*x*10^−12^, *ϵ*^2^ = 0.75, *CI*: [0.63, 1]). This quadratic trend explained the data better than assuming a linear trend (Chi-square test, *χ*^2^(3) = 46.39, *p* = 4.7*x*10^−10^).

Taken together, our behavioral results showed that participants performed the task well (Fig. 1B, C), and that their TTC estimates exhibited systematic regression-to-the-mean biases (Fig. 1B). These biases are well documented in the literature and suggest that participants relied on prior knowledge beyond the current trial to estimate time (Miyazaki et al., 2005; Jazayeri, Shadlen, 2010; Acerbi et al., 2012; Cicchini et al., 2012; Chang, Jazayeri, 2018; Meirhaeghe et al., 2021; Polti et al., 2022).

### Entorhinal cortex activity reflects trial-wise accuracy and biases in TTC estimation

Previous work on the present data showed that activity in the hippocampus reflected the magnitude of the behavioral regression-to-the-mean effect across trials (Polti et al., 2022). Here, we tested whether a similar effect can be observed for the pmEC. Using mass-univariate general linear models (GLM), we modeled the activity in each trial parametrically either as a function of accuracy (i.e., the absolute difference between estimated and target TTC; Fig. 2A, top) or as a function of the magnitude of the regression effect (i.e., the absolute difference between estimated TTC and the mean of the tested intervals; Fig. 2A, bottom). To avoid effects of potential collinearity on the final parameter estimates, these two predictors were fit in two independent GLMs, which included additional nuisance predictors such as head-motion parameters (see methods). We found that pmEC activity was higher in trials in which TTC estimates were more accurate (Fig. 2B, left; two-tailed one-sample Wilcoxon signed-rank test; *V* = 89, *p* = 1.8*x*10^−4^, *r* = −0.70, *CI*: [−0.85, −0.45]), but also that it linearly increased with stronger regression-to-the-mean biases (Fig. 2B, right; two-tailed one-sample Wilcoxon signed-rank test; *V* = 157, *p* = 0.015, *r* = −0.47, *CI*: [−0.72, −0.13]), resembling the effects previously reported for the hippocampus (Polti et al., 2022). Note that both effects were negative and showed strikingly distinct whole-brain spatial patterns (Polti et al., 2022), suggesting that they are based on independent variance in the entorhinal signal.

**Figure 2:**
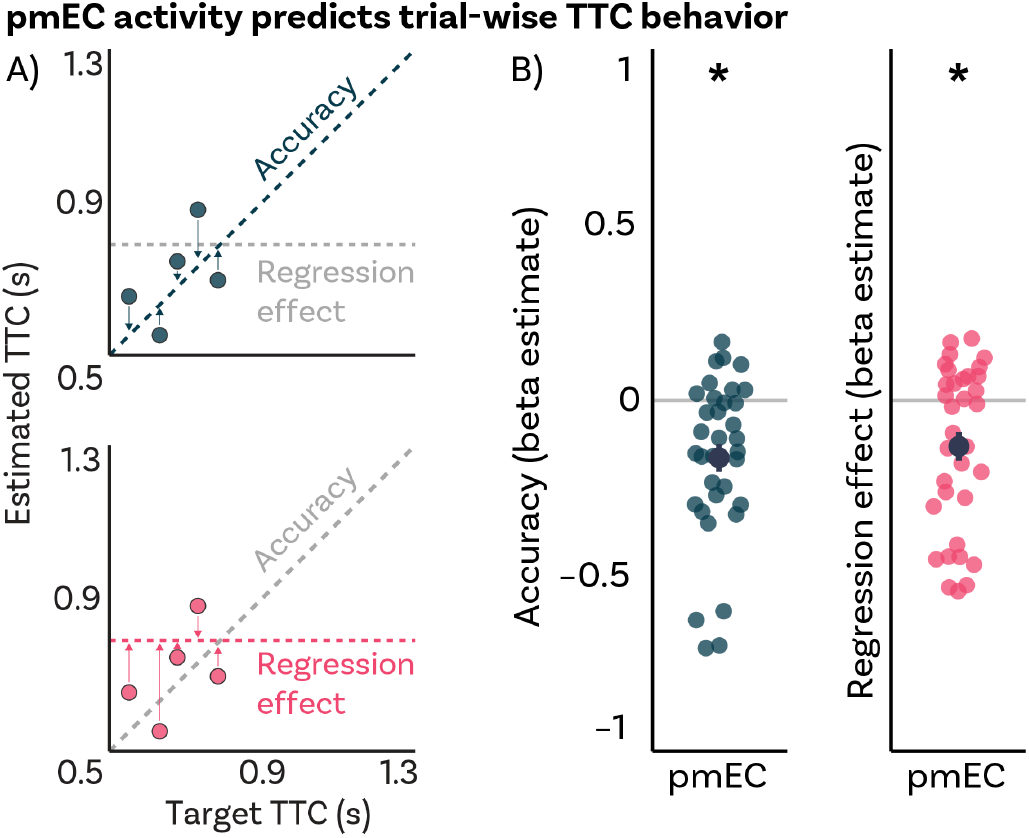
Posteromedial entorhinal cortex (pmEC) activity predicts trial-wise TTC behavior. A) Schematic description of the parametric regressor (PR) used in each separate GLM. The Accuracy PR (Top) contained the absolute difference between estimated TTC and the identity line for each trial (petrol diagonal dashed line), whereas the Regression effect PR (Bottom) contained the absolute difference between estimated TTC and mean of the tested TTCs (0.82, magenta horizontal dashed line). B) Independent regions-of-interest analysis for pmEC. We plot the beta estimates obtained for each participant for each of the two regressors. Negative values indicate higher pmEC activity with either higher accuracy (left) or higher magnitude of the regression effect (right). Depicted are the mean and SEM across participants (black dot and line) overlaid on single participant data (colored dots). Statistics reflect p<0.05 (*) obtained using a group-level two-tailed one-sample Wilcoxon signed-rank test against zero.

### Entorhinal grid-like signals predict behavioral performance in time estimation

The results reported above indicate that trial-wise pmEC activity was associated with the accuracy and the bias in TTC estimation. However, these analyses do not address whether or not grid-like signals show an association to our behavioral measures as well. Therefore, we next examined grid-like signals in our data for each TTC_*t*_ using an established quadrature filter approach (Doeller et al., 2010). The aim of this analysis was to examine whether pmEC voxels exhibited visual grid-like signals in our task (i.e., six-fold rotationally symmetric modulations as a function of gaze direction (Nau et al., 2018b; Julian et al., 2018; Staudigl et al., 2018; Graichen et al., 2025), and if so, whether these signals show a relationship to the regression-to-the-mean bias as well.

We split the time series of each pmEC voxel into two halves for later cross-validation, and then modeled each of the halves separately using a voxel-wise GLM. The GLM included two parametric main predictors modeling the sine and cosine of gaze direction modulo 60° (Fig. 3A) in addition to nuisance regressors (see methods). The ratio between the beta estimates obtained for these two predictors then allowed us to infer the putative grid orientation for each voxel (i.e., the phase of the hexadirectional modulation), which were then averaged across voxels and across scanning runs within each data partition. This procedure resulted in one putative grid-orientation for each TTC_*t*_ and each data partition. Using a second GLM, we then tested the reliability of these grid-like signals in the respective held-out data by modeling the activity of each voxel as a function of gaze direction aligned with the putative grid orientation modulo 60°. In other words, the predictor tested whether MRI signals observed for gaze directions aligned with the putative grid orientation were stronger than those observed for directions misaligned to it.

**Figure 3:**
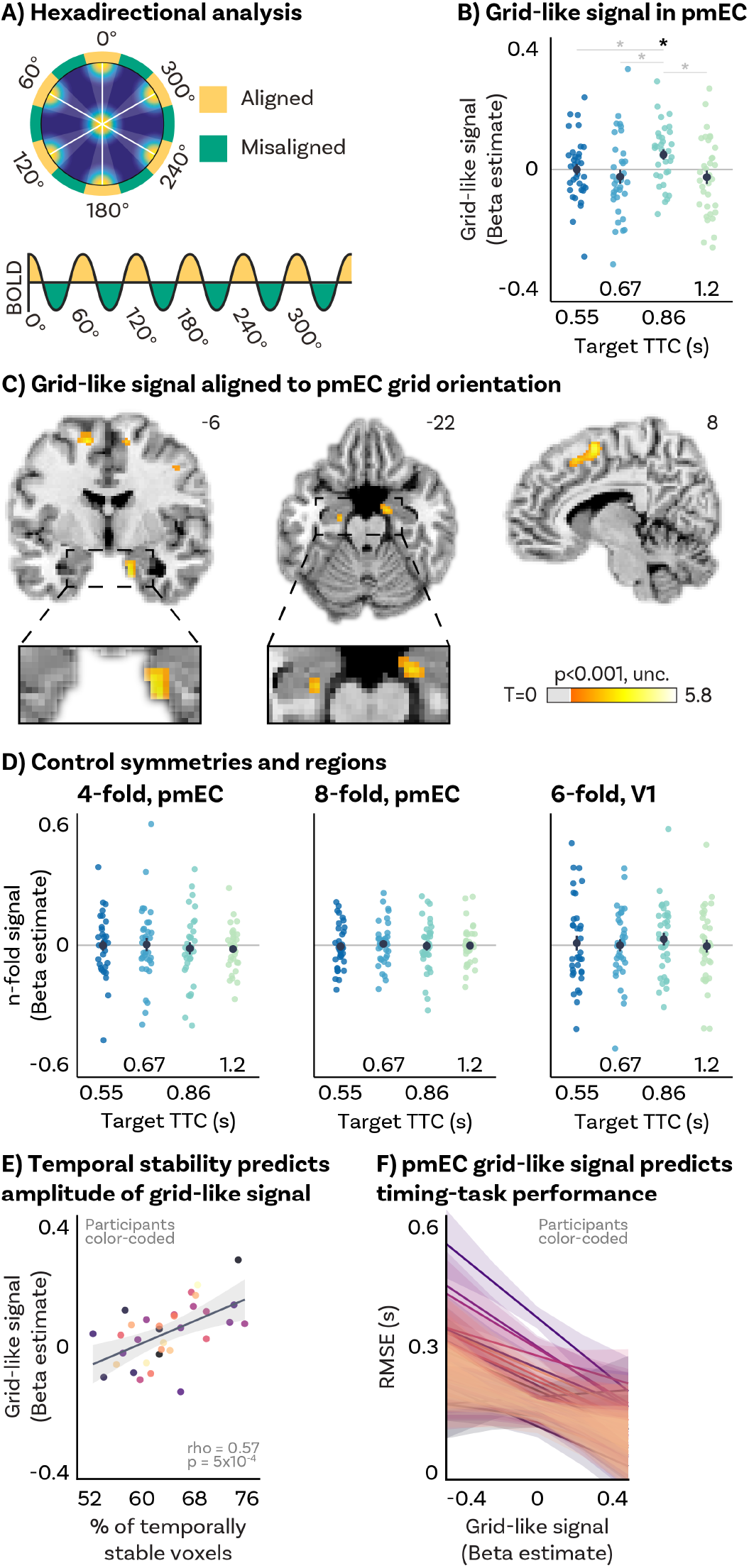
A) Visual grid-like analysis method. The hexadirectional signal is cross-validated across data partitions. Putative grid-orientation was estimated using half of the data and then used to contrast orientation-aligned vs. orientation-misaligned gaze movements in the other half (odd vs. even runs). B) Independent regions-of-interest (ROI) analysis for 6-fold symmetry in pmEC. We plot the amplitude of the hexadirectional signal in held-out data expressed as beta estimate. Positive Beta values indicate stronger grid-orientation alignment between data partitions. We found reliable cross-validated hexadirectional modulation at the group level only for TTC_0.86_ trials. There were consistent differences in pmEC fMRI activity for aligned *vs*. misaligned directions across target TTCs. Statistics reflect p<0.05 at FDR-corrected levels (*). C) Voxel-wise analysis results exhibiting activity modulation by gaze movement direction with 60° periodicity aligned to the pmEC grid orientation for TTC_0.86_ trials. We plot thresholded t-test results at 2mm resolution at p < 0.001 uncorrected levels overlaid on a structural template brain. Insert zooming in on EC and MNI coordinates added. D) Control symmetries and regions. Left: 4-fold symmetry in pmEC. Center: 8-fold symmetry in pmEC. Right: 6-fold symmetry in V1. E) TTC_0.86_ pmEC temporal stability predicts corresponding grid-like modulation across participants. Each dot represents a single participant. Regression line (black) and standard error (gray shade). F) Within-subject pmEC grid-like modulation predicts TTC estimation error (RMSE). Stronger pmEC grid-like modulation elicited lower RMSE. Separate linear mixed-effect model regression fits and standard error shading are plotted for each participant. B,D) Depicted are the mean and SEM across participants (black dot and line) overlaid on single participant data (colored dots). Target TTCs are color coded. EF) Participants are color coded.

We found that pmEC indeed exhibited a reliable grid-like modulation in our task as indicated by a regions-of-interest (ROI) analysis. Critically, however, we observed a main effect of TTC_*t*_ on the amplitude of this grid-like modulation (Fig. 3B; MEM, *F*(3) = 3.08, *p* = 0.029, *ϵ*^2^ = 0.04, *CI*: [0, 1]), with only one of the four tested intervals showing the effect (TTC_0.86_, Fig. 3B; Table S1). Trials in which TTC_0.86_ was tested yielded a stronger grid-like signal than all other TTCs (Table S1). Note that TTC_0.86_ was the interval that was closest to the mean of all intervals. A whole-brain voxel-wise analysis later confirmed that this grid-like modulation for TTC_0.86_ occurred in both hemispheres in the entorhinal cortex (Fig. 3C, left and middle panels) as well as in a few other regions that shared the putative grid orientation with the pmEC (e.g., the pre-supplementary motor area (preSMA), Fig. 3C, right panel; see Table S2 for post-hoc ROI-analysis). No effect was observed in pmEC when the same cross-validation analysis was repeated for 90° (Fig. 3D, Left; Table S3A) and 45° (Fig. 3D, Center; Table S3B), ruling out that other periodicities or single directions underly our results. Furthermore, as expected based on previous work (Nau et al., 2018b), the early visual cortex did not exhibit a grid-like signal (Fig. 3D, Right; Table S3C). Note that eye tracking-based fixation error was equivalent across target TTCs and gaze movement directions (Fig. S2; Table S8), ruling out that potential variations in grid-like signals could be attributed to differences in eye movements.

Having established that pmEC activity was modulated in a grid-like fashion for one of the TTCs, we next sought to understand the underlying differences across TTCs in more detail. Previous studies using navigation paradigms (Kunz et al., 2015; Stangl et al., 2018) suggested that these differences may be due to (i) differences in spatial stability (i.e., grid orientations may differ across voxels and therefore average out), or (ii) differences in temporal stability (i.e., grid orientation may change over the course of the experiment). To test whether any one of these factors could explain the pattern of results in our data, we estimated the spatial and temporal stability of the pmEC grid-like signal for each of the 4 TTCs separately. We found that grid orientations clustered across pmEC voxels for all TTC_*t*_ in a similar way (Fig. S4B; Table S4A), ruling out differences in spatial stability between TTC_*t*_ (Fig. S4B; MEM, *F*(3) = 0.36, *p* = 0.78, *ϵ*^2^ = −0.02, *CI*: [0, 1]). However, temporal stability predicted the amplitude of grid-like signals, meaning that the more pmEC voxels showed a stable grid orientation over data partitions, the stronger the resulting cross-validated signal amplitude turned out to be (Fig. 3E; Spearman’s *rho* = 0.57, *p* = 5*x*10^−4^). Furthermore, as expected based on the results for grid-like signal amplitude, TTC_0.86_ trials showed the highest temporal stability among all tested TTC_*t*_ (Fig. S4A; Table S4B, MEM, *F*(3) = 3.49, *p* = 0.018, *ϵ*^2^ = 0.05, *CI*: [0, 1]).

These findings provide evidence that pmEC grid-like signal amplitude differed across tested intervals, and that this difference was largely explained by how stable the putative grid orientation was over time. Importantly, we observed the grid-like signal only for the interval closest to the mean of all intervals. This result dovetails with our prior observation that trial-wise pmEC activity correlated with the regression-to-the-mean bias in behavior (Fig. 2), and suggests that grid-like signals may also show a relationship to task performance. This was indeed the case: within each participant, higher grid-like signal amplitude predicted lower RMSE, after controlling for the effect of *TTC*_*t*_ (Fig. 3F; MEM, *F*(1, 24.68) = 5.85, *p* = 0.023, *ϵ*^2^ = 0.16, *CI*: [0, 1]). Again, this result was not observed for early visual cortex (V1, Fig. S4E; MEM, *F*(1, 27.85) = 1.68, *p* = 0.205, *ϵ*^2^ = 0.02, *CI*: [0, 1]).

To understand the relationship between pmEC grid-like signals and timing behavior in more detail, we split the summary metric used in the previous analysis (RMSE) into its two main components (Variability and Bias) and tested their relationship to grid-like signals. We found that pmEC grid-like signal amplitude predicted within-participant response variability differences across TTC_*t*_ above and beyond what can be explained by scalar variability (Fig. S1B, Left panel; MEM, *F*(1, 25.35) = 7.42, *p* = 0.012, *ϵ*^2^ = 0.2, *CI*: [0.02, 1]). However, no within-participant association was found between response bias differences across TTC_*t*_ and grid-like signal amplitude (Fig. S1B, Right panel; MEM, *F*(1, 29.66) = 0.7, *p* = 0.411, *ϵ*^2^ = 0, *CI*: [0, 1]).

Finally, to asses the relationship between grid-like signals and behavioral performance irrespective of TTC_*t*_, we split trials based on the feedback participants received (reflecting high, medium or low response bias) and performed the same cross-validation analysis described above but separately for each performance level. No effect of performance level on the amplitude of the grid-like modulation (Fig. S4D; MEM, *F*(2, 99) = 1.08, *p* = 0.344, *ϵ*^2^ = 1.56*x*10^−3^, *CI*: [0, 1]), nor a reliable grid-like modulation for any of the three performance levels was found (Table S5).

### A Bayesian observer model explains performance differences across target TTCs

The behavioral and physiological results presented above suggest that participants’ duration estimates depended on both the current trial and previous trials. This pattern of results is reminiscent of prior work on Bayesian integration (Körding, Wolpert, 2004; Petzschner et al., 2015), which has been argued to underlie contextual effects in interval timing (Jazayeri, Shadlen, 2010; Acerbi et al., 2012; Cicchini et al., 2012). According to the Bayesian framework, the regression-to-the-mean effects we observed at the behavioral level (Fig. 1B) may be a natural consequence of integrating the sensory evidence in each trial (i.e., the inferred probability of a certain TTC being sampled) with an expectation informed by the statistical regularities governing prior trials (i.e., the inferred prior distribution of all sampled intervals), leading to the characteristic behavioral bias towards the mean of the encoded interval distribution. A Bayesian observer model may therefore provide a parsimonious computational explanation of both the behavioral regression-to-the-mean effect (Fig 1B) and the observed difference in grid-like activity across TTCs (Fig. 3B).

We tested this possibility post-hoc using a Bayesian observer model that was successfully used to model timing behavior in previous work (Jazayeri, Shadlen, 2010, 2015; Remington et al., 2018; Chang, Jazayeri, 2018). Briefly, the Bayes Least-Squares (BLS) model predicts the optimal posterior TTC estimate for each trial by combining two sources of information: (i) the probability that a specific TTC was sampled given the sensory evidence obtained during the visual tracking phase (Likelihood, Fig. 4A, Left), and (ii) a Gaussian prior representing the inferred distribution of sampled intervals (Prior; Fig. 4A, Left). By applying Bayes’ Theorem to compute a posterior distribution, the model identifies the mean of this distribution as the final estimate, thereby minimizing the expected squared error. To evaluate whether participants combined Prior and Likelihood sources of information when estimating the TTC, we compared the original BLS model to two alternatives that either ignore the Prior (Maximum-Likelihood Estimation, MLE model) or Likelihood (PRI model). However, these alternative models did not explain our data as well as the BLS model did (Fig. 4B; Table S6A). Additional control models confirmed that our results are better explained by a strengthening of the prior over the course of learning, rather than by changes in visual attention (Fig. S5, see Methods).

**Figure 4:**
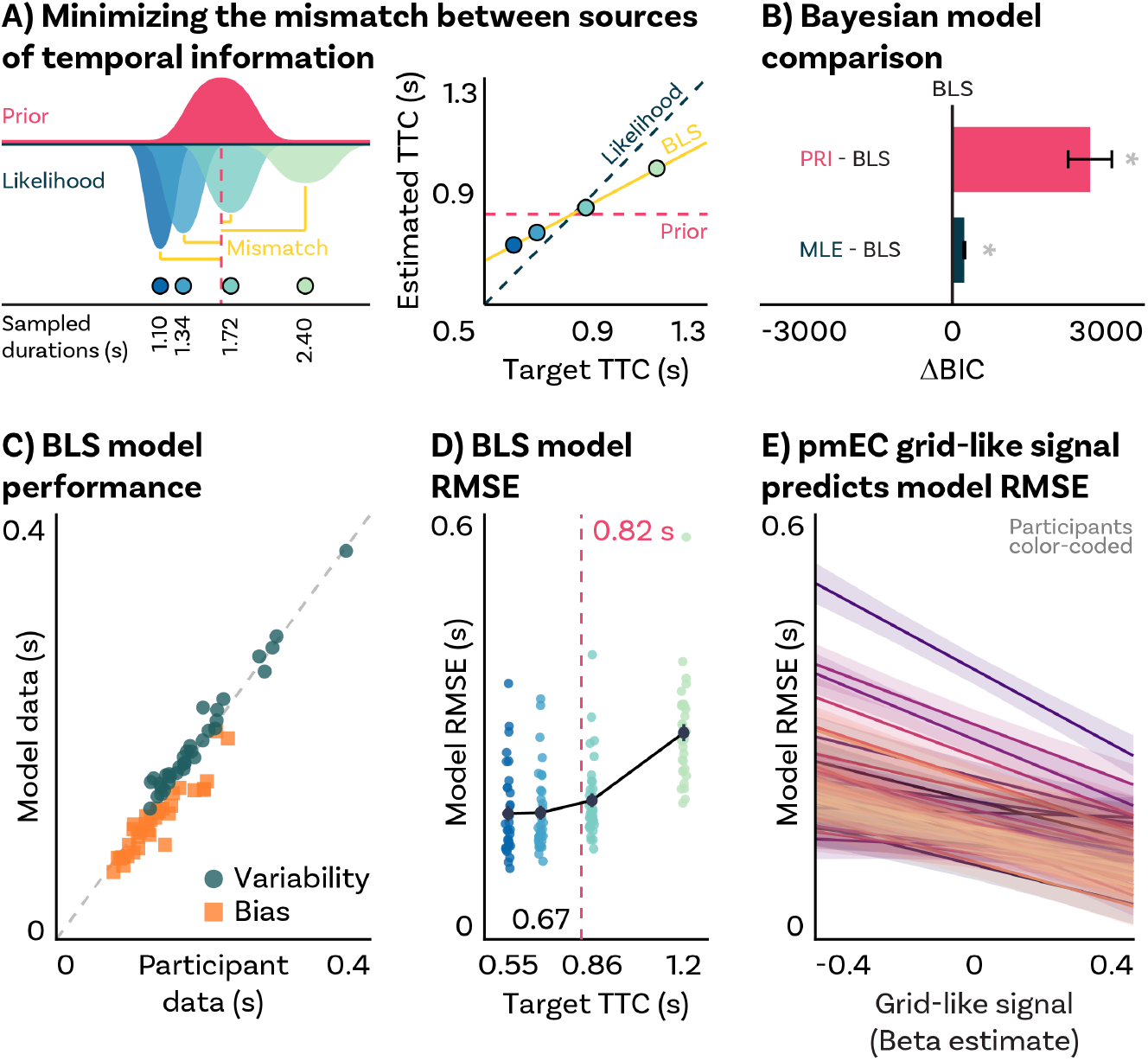
A) Schematic illustration of Bayesian TTC estimation. Left: A Bayes Least-Squares (BLS) observer integrates sensory Likelihood and Prior information sources to minimize their mismatch (yellow lines). One source of information comes from a prior expectation (Prior, magenta Gaussian distribution), centered at the mean of the sampled durations (1.64 s, magenta vertical dashed line). Another source of information comes from noisy sensory inputs (Likelihood, Gaussian distributions of different widths and turquoise tones). On each trial, by combining these two information sources participants can produce a statistically optimal TTC estimate. Right: As a result of the integration of these two sources of information, TTC estimates (colored dots over yellow bold line, BLS) are biased ‘away’ from the identity line (Likelihood, petrol blue diagonal dashed line) and ‘towards’ the mean of the sampled TTCs (Prior, magenta horizontal dashed line). B) Bayesian model comparison. The BLS model was compared to a Maximum Likelihood Estimation (MLE) model that neglects Prior information and a Prior-dominant (PRI) model that neglects Likelihood information. Each model was fit to all participants, and the Bayesian Information Criterion (BIC) was used as a statistical comparison metric. We expected the BLS model to show the lowest relative BIC value among them all, indicating the best trade-off between goodness-of-fit and complexity. We calculated the difference in BIC between each competing model and the BLS model (ΔBIC). Positive (negative) values represent evidence against (in favor) the competing models. Statistics reflect p<0.05 at FDR-corrected levels (*) from one-tailed one-sample Wilcoxon signed-rank tests against 0 (Table S6). Competing models are are color-coded. Group-level SEM depicted as error-bars. C) BLS model performance. We plot the average Variability (green circles) and Bias (orange squares) for each participant computed from 100 simulations of the BLS model vs. the same metrics computed from participants’ responses. The BLS model successfully captures participants’ responses as evidenced by the alignment with the identity line (diagonal grey dashed line). Response metrics are color-coded. D) BLS model RMSE per target TTC. We plot the average RMSE values for each participant computed from 100 simulations of the BLS model. RMSE differences across TTCs show a quadratic pattern centered on the mean of the sampled durations (0.82 s, magenta vertical dashed line), replicating real participants’ behavior (Fig. 1C). Target TTCs are color coded. Single-participant data plotted as dots. E) As expected, participants’ pmEC grid-like signals predict BLS model’s RMSE, replicating our experimental observations (Fig. 3F). Separate linear mixed-effect model regression fits and standard error shading are plotted for each participant.

From the best fitted BLS, MLE and PRI models, we then simulated each participant’s behavior 100 times and calculated the average response Variability, Bias and RMSE. In line with previous work (Jazayeri, Shadlen, 2010, 2015; Remington et al., 2018; Chang, Jazayeri, 2018), we found that the BLS model accurately captured participants’ response Variability and Bias in the TTC task (Fig. 4C), and did so better than the MLE and PRI models (Table S6B, C). Participants’ RMSE quadratic pattern across TTC_*t*_ (Fig. 1C) was also replicated by model-derived RMSE values (Fig. 4D; MEM, *F*(2, 33) = 197.39, *p* = 2.2*x*10^−16^, *ϵ*^2^ = 0.92, *CI*: [0.87, 1]). Moreover, by relating the model estimates to the neuroimaging results obtained for pmEC, we found that model-derived RMSE values were correlated with the amplitude of grid-like signals observed within participants (Fig. 3E; MEM, *F*(1, 32.48) = 6.49, *p* = 0.016, *ϵ*^2^ = 0.14, *CI*: [0.01, 1]). Together, these modeling results suggest that the differences in participants’ timing behavior and in grid-like signals across TTC_*t*_ can indeed be well explained using a Bayesian observer model that assumes that predictions informed by temporal regularities bias timing behavior.

## Discussion

The present study examined whether and how the human pmEC contributes to the encoding of task regularities that guide timing behavior. We used fMRI to record human brain activity while participants performed a rapid time-to-contact estimation task, which allowed us to analyze trial-wise pmEC activity as a function of time-estimation performance across a range of sampled intervals. Moreover, the task included periods in which participants followed a moving fixation target with their eyes, allowing us to estimate visual grid-like signals in the pmEC: the amplitude and stability of a six-fold rotationally symmetric MRI-signal modulation as a function of gaze direction. Note that this signal was specifically six-fold rotationally symmetric and not driven by individual movement direction, as an analysis of control symmetries showed (Fig. 3D). We found that pmEC activity closely tracked task performance across trials as well as behavioral response biases towards the mean interval, and we observed a bias towards that mean interval on the level of grid-like signals. Traditionally, such regression-to-the-mean biases have been taken as evidence for Bayesian integration in the brain (Petzschner et al., 2015), since they are well explained by models that assume the integration between sensory evidence with a prior expectation derived from learned statistical regularities. Indeed, a Bayesian observer model previously used to model timing behavior in humans and macaques (Jazayeri, Shadlen, 2010, 2015; Remington et al., 2018; Chang, Jazayeri, 2018; De Kock et al., 2021) provided a parsimonious account for both our behavioral and pmEC results. In the following, we will discuss these results in light of previous literature on timing behavior and relate them to prior work on spatial and non-spatial coding principles in the hippocampal formation.

### Task regularities bias TTC estimation

While participants were not explicitly told about the true range of intervals that were tested, their estimates were nevertheless biased towards the mean interval (Fig. 1B, C). This regression-to-the-mean effect in time estimation is well documented and has been proposed to reflect participants’ reliance on the temporal regularities learned from previous trials (Miyazaki et al., 2005; Jazayeri, Shadlen, 2010; Acerbi et al., 2012; Cicchini et al., 2012; Petzschner et al., 2015; Chang, Jazayeri, 2018; Polti et al., 2022). Specifically, integrating a prior expectation with the sensory evidence obtained in the current trial may help participants to anticipate the trajectory of moving objects during occlusion (Fig. 1A). This prior expectation likely takes the form of a Gaussian distribution of time intervals centered on the mean (Fig. 4A). Relying on this prior to make an interval judgment will therefore most often be biased towards that mean, depending on how strong the sensory evidence is in a given trial about the true TTC that was tested. With increased mismatch between prior expectation and sensory evidence, participants’ estimates may be biased more towards the mean (Fig. 1B, C), potentially reflecting an increase in uncertainty about their estimate (Jazayeri, Shadlen, 2010; Petzschner et al., 2015). Overall, this strategy may lead to large errors in some trials, but it may nevertheless be adaptive, since the mean is still a “good guess” for the large majority of trials. By reducing response variability (Fig. S5A), these biases may especially be useful when evidence derived from the senses is sparse or noisy. While we tested only four intervals drawn from a uniform distribution, previous work has shown that interval estimates tend to be encoded using a Gaussian distribution even when the intervals were sampled uniformly (Acerbi et al., 2012; Hahn, Wei, 2024). This is in line with our observation that our data was well described using a Bayesian model that assumed a Gaussian distribution centered on the mean interval (Fig. 4C).

### Entorhinal cortex encodes task regularities that afford time estimation

Entorhinal activity reflected the behavioral response biases towards the mean across trials, as well as overall task performance. In our view, this result is striking as the entorhinal cortex is mostly studied in the context of spatial navigation (Epstein et al., 2017; Kunz et al., 2019) and long-term memory formation (Fernández et al., 1999; Hargreaves et al., 2012; Staresina et al., 2013; Schiller et al., 2015), not in the context of rapid timing tasks. Notably, similar tasks have been used successfully for studying predictive processes in rodents (Henke et al., 2021), humans (Jazayeri, Shadlen, 2010; Acerbi et al., 2012; Cicchini et al., 2012; Chang, Jazayeri, 2018; Polti et al., 2022), and non-human primates (Jazayeri, Shadlen, 2015; Wang et al., 2018; Sohn et al., 2019), but not in the context of entorhinal functions. Our results suggest that human entorhinal cortex contributes to timing task performance in real time as the task is being performed. In fact, we found trial-wise pmEC activity to follow a pattern that closely resembled the one previously reported for the adjacent hippocampus, as well as for other regions prominently including the striatum (Polti et al., 2022; Rolando et al., 2024). The contributions of the entorhinal cortex to task performance may therefore closely depend on other regions shown to preferentially encode distinct behaviorally relevant factors, such as task details versus task structure (Doeller et al., 2008; Geerts et al., 2020). Specifically, the hippocampal-entorhinal region has been shown to signal the encoding of task structure (e.g., graphs reflecting transition probabilities between sequentially presented stimuli (Garvert et al., 2017) and abstract task spaces (Constantinescu et al., 2016; Theves et al., 2019, 2020; Viganò, Piazza, 2020; Park et al., 2021), which may reflect the learning of generalizable principles that guide behavior across tasks. Our results are in line with these ideas and support recently proposed computational accounts of entorhinal function that center on structured, factorized representations that afford inference and generalisation (Whittington et al., 2020). Based on these accounts, as well as prior work showing that hippocampal activity tracks the learning of TTC regularities in our data (Polti et al., 2022), we speculate that our results reflect a division of labor in which the hippocampus updates a temporal prior while pmEC grid-like signals encode this learned task structure for ongoing time estimation.

Our results further dovetail with work on temporal-context encoding (Schapiro et al., 2012; Hsieh et al., 2014) and sequence memory (Fortin et al., 2002; Montchal et al., 2019; Bellmund et al., 2020a, 2022) in the hippocampal-entorhinal region. For example, animal studies have shown that damage to the entorhinal cortex impairs memory for relations (Buckmaster et al., 2004), and inactivation of the rodent MEC prevents context-dependent learning of intervals (Bigus et al., 2024). In general, recent years have seen a growth in the literature on the links between MEC and timing behavior in rodents (Heys, Dombeck, 2018; Heys et al., 2020; Vo et al., 2021; Dias et al., 2021; Bigus et al., 2024), and showing that entorhinal activity and whole-brain functional connectivity reflects the passage of time in humans (Wang et al., 2025). In addition, time-dependent activity has been demonstrated in the neighboring lateral EC of rodents (Tsao et al., 2018; Kanter et al., 2025) and in its human homologue (Montchal et al., 2019; Bellmund et al., 2019). Our results extend these reports to the human pmEC, revealing a direct relationship between pmEC activity and behavioral performance in a timing task. Specifically, its activity mirrored a behavioral bias towards the mean of the tested intervals; a phenomenon that occurs for any type of magnitude estimation (Petzschner et al., 2015; Petzschner, Glasauer, 2011). Investigating the relationship between these mean biases and neural activity across a range of tasks therefore provides fertile ground for investigations of the domain-general functions of the hippocampal-entorhinal region (Behrens et al., 2018; Bellmund et al., 2018; Stachenfeld et al., 2017). In line with this idea, hippocampal activity has been linked to regression-to-the-mean biases during the estimation of spatial distances (Wiener et al., 2016) and intervals (Polti et al., 2022), and entorhinal activity has been shown to be modulated by spatial context during virtual navigation (Julian, Doeller, 2021).

### Non-spatial task factors shape entorhinal grid-like signals

In addition to co-variations in pmEC activity and task-performance measures across trials, we found that pmEC grid-like signals in particular reflected participants’ timing performance across intervals (Fig. 3F). Specifically, within-participant differences in timing variability were associated with differences in grid-like signal magnitude (S1B, left). Like behavioral performance, also the grid-like signal seemed to be biased towards the mean interval, with cross-validation revealing a robust grid-like modulation exclusively for the interval closest to the mean (Fig. 3B). This effect was driven by the temporal stability, not spatial stability, of the grid-like signal (Figs. S4A, B), in line with previous reports (Kunz et al., 2015; Stangl et al., 2018). Furthermore, our results also replicate previous work on visual grid-like signals in humans (Nau et al., 2018b; Julian et al., 2018; Staudigl et al., 2018; Graichen et al., 2025), while going beyond these studies by reporting a relationship between these signals and timing performance. Eye movements may engage similar processes in the entorhinal cortex as navigation (Nau et al., 2018a), while offering higher experimental control and study-design efficiency. Since behavioral performance was well explained by a Bayesian observer model, we tested whether this model also predicted the grid-like signal differences across intervals, which was indeed the case (Fig. 4E). Model simulations recapitulated the link between task performance and the grid-like signal differences across intervals, potentially offering a parsimonious and normative explanation of our findings. Furthermore, the modeling results suggest that participants’ performance gains were likely driven by learning of the temporal context, decreasing response variability while increasing the bias towards the mean (Fig. S5). The entorhinal cortex, and grid cells in particular, may support the encoding of task regularities that ultimately manifest in both neural activity and behavior as regression-to-the-mean effects.

Why would a spatial grid-like signal be modulated by the range of intervals that was tested? Previous work has shown that task-dependent factors such as environmental features, goal locations, and rewards can distort the grid-cell pattern in rodents (Barry et al., 2007, 2012; Krupic et al., 2015; Hardcastle et al., 2015; Keinath et al., 2018; Boccara et al., 2019; Butler et al., 2019), as well as grid-like (Viganò et al., 2023) and behavioral (Bellmund et al., 2020b; Chen et al., 2015) response patterns in humans. Notably, grid cell distortions in rats correlate with distance estimation errors (Duncan et al., 2025). It therefore seems plausible that also human visual grid-like signals can be modulated by behaviorally-relevant features of the task and correlate with participants’ TTC estimation error. In our task, the most relevant features were the intervals that were tested, possibly explaining why the timing bias was lowest (Fig. S1B, right) and the temporal stability of the grid-like signal was highest (Fig. S4A) for the interval closest to the mean. Previous work suggested that the distortions of the grid-cell patterns could be the consequence of conflicting sources of information (Barry et al., 2007; Hardcastle et al., 2015; Krupic et al., 2015), which would be broadly in line with our Bayesian modeling results (Fig. 4). Because spatial information remains constant across trials, in our task the conflicting sources of information may be the temporal prior expectation derived from previous trials, which is inherently biased towards the mean interval, and the current-trial evidence of a sampled duration derived from the senses. Therefore, we speculate that the amplitude and orientation of grid-like signals depend on the agreement between these two sources of information. This alignment becomes higher the closer the tested interval is to the mean interval.

### Predictive processing as a domain-general principle of entorhinal function?

While many of the above considerations remain to be tested, it has long been recognized that learning task regularities requires the encoding of relationships between stimuli, actions, and events (Körding, Wolpert, 2004; Petzschner, Glasauer, 2011; Petzschner et al., 2015), which has been proposed to build on a relational coding scheme that has often been discussed for the medial temporal lobe (Manns, Eichenbaum, 2006; Whittington et al., 2020) even for non-spatial feature spaces (e.g., Aronov et al. (2017); Bellmund et al. (2018); Behrens et al. (2018); Constantinescu et al. (2016); Bao et al. (2019); Theves et al. (2019, 2020); Viganò et al. (2021); Park et al. (2021); Wagner et al. (2023); Nitsch et al. (2023)). Notably, non-spatial entorhinal coding also includes grid-like representations during navigation through an abstract age–by–time-of-day space (Peters-Founshtein et al., 2024), which differs from the present findings. In contrast to this previous report, our results indicate that task-relevant stimulus features directly shape entorhinal spatial coding. Our task explicitly required participants to make temporal predictions, but the central ideas and observations presented in this article may therefore very well translate also to other tasks and less explicit situations. Taking a predictive-processing perspective may more generally help to understand the functional contributions of the entorhinal cortex across behavioral domains and species (e.g., Cothi de et al. (2022)), which includes, but is not limited to, the Bayesian framework. In fact, a growing number of studies support the idea that Bayesian models can provide a normative explanation for a range of observations in the spatial navigation literature, including the integration of visual cues and landmarks during path integration (Cheng et al., 2007; Petzschner, Glasauer, 2011; Kessler et al., 2022). Intriguingly, the same Bayesian model (Kang et al., 2023) that explains distortions in spatial memory in humans (Hartley et al., 2004; Chen et al., 2015; Bellmund et al., 2020b; Keinath et al., 2021) can also explain distortions of the grid-cell pattern in rodents (e.g., Krupic et al. (2015)), pointing towards a unified computational account for behavior and entorhinal activity across species.

### Predictions for future work

Our results suggest that non-spatial task factors shape human entorhinal activity in real time as the task is performed, a phenomenon that future studies should investigate across a range of tasks. It is important to note that we modeled our results using a Bayesian observer model due to its successful application to similar problems in the past (Jazayeri, Shadlen, 2010; Acerbi et al., 2012; Remington et al., 2018; Chang, Jazayeri, 2018), but other computational frameworks may be able to explain our data equally well. Approaches that model the mismatch between prior expectations and sensory evidence more explicitly seem especially promising in this context (e.g., prediction errors in reinforcement learning (Momennejad et al., 2017; Niv, 2009; Dayan, Daw, 2008; Stachenfeld et al., 2017)). Second, our results are in line with the idea that participants encoded the tested intervals using a Gaussian distribution centered on the mean intervals, but a more fine-grained sampling of intervals would be necessary to convincingly show that this was actually the case. The ideal scenario would be to sample many more time intervals, not from one but from multiple distributions, each centered on a different value. In this case, one would predict that the grid-like signal is stable at the center of each of these distributions. Another interesting avenue would be to examine how grid cell-like signals develop as task regularities are learned. However, this would require a new analytical approach capable of estimating this signal within each scanning run individually, unlike current approaches that rely on across-session cross-validation. Finally, to investigate the range of non-spatial factors that affect grid-like signals, and therefore to understand potential domain-general contributions of the pmEC to human cognition, future work should test a range of tasks beyond time estimation and spatial navigation (Nau et al., 2024).

## Conclusion

Taken together, using fMRI and a rapid time-to-contact estimation paradigm in humans, we showed that time estimates are biased towards the mean of the tested intervals, and that this mean bias is reflected in entorhinal activity across trials (similar to the hippocampus, Polti et al. (2022)). Moreover, we report a novel relationship between grid-like signals and behavioral performance, as the amplitude of the grid-like signal correlated with participants’ time-estimation error, and the putative grid orientation was stable exclusively for the interval closest to the mean. Finally, both our behavioral and neuroimaging results were well explained (post hoc) by a Bayesian observer model that assumes the integration of prior expectations and sensory evidence. These results point to an involvement of the pmEC in interval timing, likely building on the encoding of task regularities that afford predictive inference in the service of goal-directed behavior.

## Acknowledgements

This work is funded by the European Research Council (ERC-CoG GEOCOG 724836 awarded to CFD). CFD’s research is further supported by the Max Planck Society, the Kavli Foundation, the Jebsen foundation, the Centre of Excellence scheme of the Research Council of Norway – Centre for Neural Computation (223262/F50), The Egil and Pauline Braathen and Fred Kavli Centre for Cortical Microcircuits and the National Infrastructure scheme of the Research Council of Norway – NOR-BRAIN (197467/F50). RK’s research is supported by a CIDEGENT grant (CIDEGENT/2021/027) from the Valencian Community’s program for the support of talented researchers, and by a Ministerio de Ciencia, Innovación y Universidades grant (PID2021-122338NA-I00). M.N. is supported by a Feodor Lynen Research Fellowship provided by the Alexander von Humboldt Foundation.

## Author Contributions

MN, IP and CFD conceived the study. IP and MN designed the experimental paradigm, visualized the results, and embedded them in the literature with help from RK, VW and CFD. IP implemented the experimental code and acquired and analyzed the data with close supervision from MN. MN and IP wrote the manuscript with critical feedback from RK, VW, and CFD. CFD provided general supervision of the project and secured funding. IP and MN are shared-first authors.

## Data and code availability

Source data and analysis code are available on Open Science Framework (https://osf.io/cs8d6/), along-side pre-processed eye-tracker data (https://osf.io/mrhk9/). Raw fMRI data is available at the following G-Node Infrastructure repository: https://gin.g-node.org/ipolti/TTC_HPCF.git. The original Bayesian Observer Model has been shared by Jazayeri and colleagues and can be found here: https://jazlab.org/resources/.

## Methods

### Participants

The data used in this study were used in a previous report (Polti et al., 2022). Thirty-nine participants (16 women, 23 men, 19-35 years old) were recruited for this study. Five participants were excluded: one was excluded because the eye-tracker calibration failed, one was excluded because they did not follow the task instructions, and three participants were excluded because of technical issues during scanning. A total of 34 participants entered the analysis. The study was approved by the regional committee for medical and health research ethics (project number 2017/969) in Norway and participants gave their written consent prior to scanning in accordance with the Declaration of Helsinki (World Medical Association, 2013).

### Task

We used a Time-to-contact (TTC) task that required participants to estimate the time when a moving dot reached a target location. Each trial began with the smooth visual pursuit of a dot moving in 1 of 24 predefined linear trajectories with 1 of 4 possible speeds: 4.2°/s, 5.8°/s, 7.5°/s and 9.1°/s. All trajectories had two segments, one where the dot was visible (“Gaze trajectory”, Fig. 1A; length of 10 dva) and one where the dot was occluded (length of 5 dva). Because all trajectories had the same length, the 4 speeds led to 4 target TTC (*TTC*_*t*_) durations: 1.2s, 0.88s, 0.67s, and 0.55s. A *TTC*_*t*_ duration was defined as the time it takes the dot to traverse the occluded segment. When the dot reached the end of the visible segment, a fixation cross remained in place until participants had performed a TTC estimation judgment. The latter consisted of clicking a button when the dot presumably reached the end of the occluded segment (“Boundary”, Fig. 1A). After giving a response, participants received visual feedback for 1 s reflecting their TTC estimation accuracy. When accuracy was not high, a visual cue signaled the TTC error direction: either responding too early (underestimation) or too late (overestimation). At the feedback offset, the dot became visible again and remained static for a variable ITI sampled randomly from a uniform distribution ranging from 0.5 to 1.5 s. Then, a new trial began with the dot moving in a different direction. Over the course of 768 trials, we sampled eye movement directions with 15° resolution. Participants were never explicitly informed about the full visual trajectory or the range of *TTC*_*t*_. See previous work of Polti et al. (2022) for more details on the task.

### Behavioral analysis

Participants indicated the estimated TTC (TTC_*e*_) in each trial via button press. To test if participants discriminated the four target TTCs we used a linear mixed-effect model with TTC_*e*_ as the dependent variable, target TTC (TTC_*t*_) as the predictor and separate intercepts and TTC_*t*_ slopes per participant:

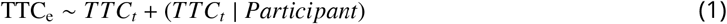

We partitioned participants’ behavioral performance for each target duration TTC_*t, j*_ (where *j* ∈ {0.55, 0.67, 0.86, 1.2}) using two metrics, Variability and Bias (Fig. S1A). Variability _*j*_ was computed as the standard deviation of the TTC_*e, j*_:

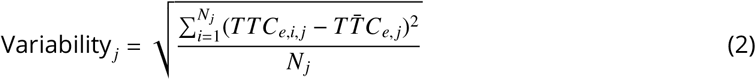

Bias _*j*_ was computed as the absolute difference between the average 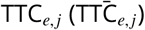 and TTC_*t, j*_:

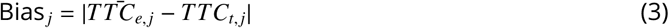

These two parameters can be integrated in a single metric that reflects the Variability-Bias trade-off when written as a Pythagorean sum, i.e. the root-mean-square error (RMSE):

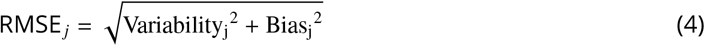

In order to analyze the pattern of RMSE across target TTCs, we used a mixed-effect model with RMSE as the dependent variable, TTC_*t*_ as a second-order orthogonal polynomial, and separate intercepts and TTC_*t*_ slopes per participant (Fig. 1C).

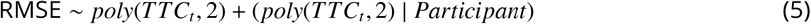

We also ran the same model again while using a linear instead of quadratic trend. We then tested which model best explained the changes in RMSE across target TTCs using a chi-square test.

### Imaging data acquisition & preprocessing

Imaging data were acquired on a Siemens 3T MAGNETOM Skyra located at the St. Olavs Hospital in Trondheim, Norway. A T1-weighted structural scan was acquired with 1mm isotropic voxel size. Following EPI-parameters were used: voxel size=2mm isotropic, TR=1020ms, TE=34.6ms, flip angle = 55°, multiband factor=6. Participants performed a total of four scanning runs of 16-18 minutes each including a short break in the middle of each run. Functional images were corrected for head motion and co-registered to each individual’s structural scan using SPM12 (www.fil.ion.ucl.ac.uk/spm/). We used the FSL topup function to correct field distortions based on one image acquired with inverted phase-encoding direction (https://fsl.fmrib.ox.ac.uk/fsl/fslwiki/topup). Functional images were then spatially normalized to the Montreal Neurological Institute (MNI) brain template and smoothed with a Gaussian kernel with full-width-at-half-maximum of 4 mm for regions-of-interest analysis or with 8 mm for whole-brain analysis. Time series were high-pass filtered with a 128 s cut-off period.

### Region-of-interest (ROI) definition and analysis

Rodent studies have consistently reported grid cells in the medial entorhinal cortex (Heys et al., 2014; Moser et al., 2014), which likely corresponds to the posteromedial portion of the entorhinal cortex in humans (Maass et al., 2015; Navarro Schröder et al., 2015). We therefore used a (bilateral) posteromedial entorhinal cortex (pmEC) mask from Navarro Schröder et al. (2015) in all our fMRI analyses (Fig. S3). As a control region, we chose the early visual cortex (V1) for which, to our knowledge, no hexadirectional signals have been reported. V1 masks were generated for each individual participant based on the automatic parcellation derived from FreeSurfer’s structural reconstruction (https://surfer.nmr.mgh.harvard.edu/). A pre-supplementary motor area (preSMA) mask was obtained from the JuBrain SPM anatomy toolbox (https://www.fz-juelich.de/inm/inm-1/EN/Forschung/_docs/SPMAnatomyToolbox/SPMAnatomyToolbox_node.html) in order to post-hoc confirm whether activity observed in voxel-wise analyses corresponded to preSMA. All masks were spatially normalized to the MNI brain template space and re-sliced to the functional imaging resolution using SPM12.

All ROI analyses described in the following were conducted using the following procedure. We extracted beta weights estimated for the respective regressors of interest for all voxels within a region in both hemispheres, averaged them across voxels within that region and performed a one-sample Wilcoxon test on group level against zero as implemented in the software R (https://www.R-project.org).

### EC activity as a function of accuracy and as a function of the regression effect

To examine the relationship between behavioral biases and brain activity, we used mass-univariate general linear models (GLM) to model the trial-wise activity of the pmEC voxels as a function of accuracy (i.e. the absolute difference between estimated and target TTC in each trial) and as a function of the regression effect (i.e. the absolute difference between the estimated TTC and the mean of the tested intervals, which was 0.82 s) in the TTC task. To avoid effects of potential co-linearity between these regressors, we estimated model weights using two independent GLMs, which modeled the time course of each trial with either one of the two regressors. The GLMs also included one regressor modeling ITIs, one for button presses and one for periods of rest, which were all convolved with the canonical hemodynamic response function in SPM12. In addition, the models included the six realignment parameters obtained during pre-processing as well as a constant term modeling the mean of the time series. After fitting each model, we used the weights estimated for the two regressors to perform ROI analyses for the EC using a two-tailed one-sample Wilcoxon test (Fig. 2B).

### Hexadirectional analysis of visual grid-like representations

Prior work showed that the MRI signal in the human entorhinal cortex exhibits a six-fold rotationally symmetric modulation as a function of gaze direction, which is thought to reflect grid-cell population activity (Nau et al., 2018b; Julian et al., 2018). Here, we tested whether such grid-like signals were also detectable in our data. The analysis builds on cross-validation to first estimate the putative grid orientation relative to the screen, and then testing in held-out data whether gaze directions aligned to this putative grid orientation are associated with stronger MRI signals than directions misaligned to it.

To estimate the putative grid orientation, we first modeled the activity in each voxel in half of the data using a GLM (odd vs even runs) that incorporated two main regressors of interest. These regressors modeled the sine and cosine of the movement direction of the fixation target *θ* with a periodicity of 60°, i.e. sin(6 *θ*) and cos(6 *θ*). For each trial, *θ* included the tracking phase as well as the TTC estimation task phases (i.e., the time the target’s movement was invisible). In addition, nuisance regressors modeled ITIs, the feedback phase, button presses and periods of rest, and two parametric regressors modeled the effect of feedback on the activity during the feedback phase and another one modeling the effect of TTC bias on button presses. All regressors were convolved with the HRF. Because different fixation-target speeds also led to different TTCs and thus trial durations, and because the amplitude of the HRF-convolved signal scaled with duration, we re-scaled the resulting main regressors to obtain a balanced regressor amplitude for all speeds. Moreover, we added the realignment parameters and a constant term to the model. Each speed level was modeled separately. Weights for all regressors were estimated using SPM12. Note that this across-session cross-validation procedure that is crucial to our analysis precludes trials-wise analysis of grid cell-like signals.

We used the beta weights corresponding to the sine (*β* sin) and cosine (*β* cos) of each target TTC level to estimate the putative grid orientation relative to the screen *ϕ* for voxels within the entorhinal cortex using the following formula:

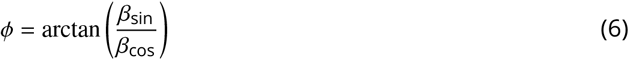

The estimated orientations were then averaged across voxels and across runs within each data partition (circular mean), resulting in a single grid orientation for each target TTC and data partition. We then used the estimated grid orientation in a second GLM to estimate the amplitude of the grid-like signal in its independent data counterpart (Fig. 3A). To do so, we again modeled nuisance variance as described before, this time adding one main regressor per target TTC modeling the cosine of each fixation-target movement direction modulo 60° aligned to the previously estimated mean orientation using the following formula:

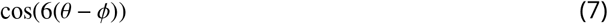

Again, all regressors but the realignment parameters and the constant term were convolved with the HRF, and all main regressors were rescaled to match their amplitude across TTCs. We then again estimated weights for all regressors using SPM12 and averaged them across the pmEC ROI. For each target TTC, we tested its corresponding estimated weight against zero using a one-tailed one-sample Wilcoxon test on group-level (Fig. 3B; Table S1). In order to test for a main effect of target TTC on grid-like signal amplitude, we ran a mixed-effects model with the estimated weights (GLS_*pmEC*_) as the dependent variable, TTC_*t*_ as predictor and participants as the error term:

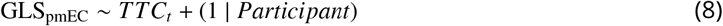

We used two-tailed paired Wilcoxon signed-rank tests to compare differences in grid-like signal between target TTCs (Table S1).

To test whether the grid-like signal found in pmEC exhibited specifically a 6-fold symmetric periodicity (60°), and not other periodicities, we repeated the cross-validation analysis for 4-fold (i.e. 90°, Fig. 3D, Left) and 8-fold (i.e. 45°, Fig. 3D, Center) symmetries. In addition, we tested for 6-fold symmetry in a control region (early visual cortex, V1; Fig. 3D, Right).

To asses the behavioral relevance of the pmEC grid-like signals after controlling for the effect of TTC_*t*_ (Fig. 3F), we ran a linear mixed effect model with RMSE as the dependent variable, with GLS_*pmEC*_, TTC_*t*_ and their interaction as predictors. The model included separate intercepts and GLS_*pmEC*_ slopes for each participant:

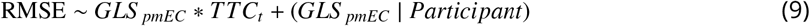

As a control, we ran the same linear mixed effect model as described above, but using instead the grid-like signals estimated in V1 (GLS_*V*1_) separately for each TTC_*t*_ (Fig. S4E):

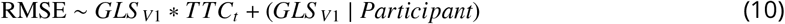

Furthermore, to analyze in more detail the relationship between pmEC grid-like signals and timing behavior we ran two separate linear mixed effect models (Fig. S1B). Each of these models shared the same predictors (GLS_*pmEC*_, TTC_*t*_ and their interaction), but differed in their dependent variable (either response Variability or Bias). Both models included separate intercepts and GLS_*pmEC*_ slopes for each participant:

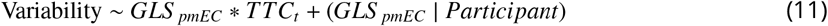

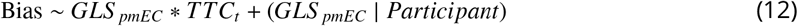

Last, we assesed the relationship between grid-like signals and behavioral performance irrespective of TTC_*t*_ by running a new hexadirectional analysis (Fig. S4D). The GLMs and cross-validation procedures were identical to the ones described before, with the only difference that instead of grouping trials by target TTC, the new hexadirectional analysis grouped trials based on the feedback participants received (reflecting high, medium or low response bias).

### Spatial and temporal stability of grid-like signals

In order to exhibit a reliable grid-like signal, a voxel’s grid orientation should remain stable over time (i.e. temporal stability). For each participant and target TTC, we therefore computed the percentage of pmEC voxels that maintained an orientation difference of less than 15° between training and test data partitions. We then tested if differences in temporal stability could explain individual differences in the pmEC grid-like modulation for TTC_0.86_ using a Spearman’s rank-order correlation (Fig. 3E).

Besides having high temporal stability, a robust grid-like signal is also expected to have high spatial stability (i.e., pmEC voxel-wise grid orientations should cluster in order to provide a robust mean grid orientation). We used Rayleigh’s *z*-value as a measure of spatial stability: higher *z* values correspond to higher voxel-wise grid orientation clustering. For each participant and target TTC (TTC_*t*_), we computed Rayleigh’s test for non-uniformity of circular data on the pmEC voxel-wise grid orientations from the training data partition. We tested the Rayleigh’s *z*-values separately for each TTC_*t*_ against zero using one-tailed one-sample Wilcoxon tests on group-level (Table S4A). In order to compare spatial stability between TTC_*t*_, we again used a mixed-effects model incorporating spatial stability as a dependent variable, TTC_*t*_ as a predictor, and participants as the error term (Fig. S4B):

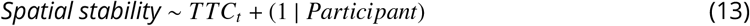

We further used a Spearman’s rank-order correlation to test if individual differences in spatial stability could predict the cross-validated grid-like signal amplitude estimated for TTC_0.86_ (Fig. S4C; Spearman’s *rho* = 0.49, *p* = 0.003).

Differences in grid-like signal amplitude across TTC_*t*_ could also be explained by differences in temporal stability. We thus used a mixed-effects model with temporal stability as a dependent variable, TTC_*t*_ as a predictor and participants as the error term to test this possibility (Fig. S4A):

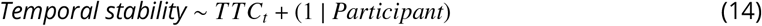

Based on the cross-validation results (Fig. 3B), we expected TTC_0.86_ to show a higher temporal stability than the other TTCs. We tested this by running one-tailed paired Wilcoxon signed-rank tests between the TTC_*t*_ (Table S4B).

### Bayesian observer model

The Bayesian observer model, developed originally by Jazayeri and colleagues (Jazayeri, Shadlen, 2010), has been successfully adapted and applied several times to model timing behavior (Jazayeri, Shadlen, 2015; Remington et al., 2018; Chang, Jazayeri, 2018; De Kock et al., 2021). We adapted the original code provided at https://jazlab.org/resources/ in order to model participants’ behavior in our TTC estimation task.

The Bayesian observer model is composed of three stages:

*Stage 1 (Measurement):* The observer makes a noisy duration measurement (*VT*_*m*_) of the visual tracking period (VT, Fig. 1A), from the movement onset of the fixation disc until it becomes occluded. The measurement *VT*_*m*_ is drawn from a Gaussian distribution centered at the true VT duration with standard deviation *σ*_*m*_ = *w*_*m*_ × *VT*, where *w*_*m*_ is the Weber fraction for VT duration measurement. This defines the likelihood function exhibiting scalar variability (Gibbon, 1977):

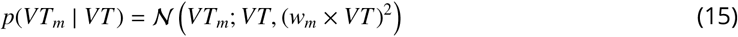

*Stage 2 (Bayesian inference):* The observer combines the measurement likelihood *p(VT*_*m*_|*VT)* with a prior distribution *p(VT)* using Bayes’ rule:

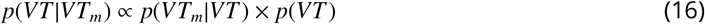

The prior *p(VT)* is as a Gaussian distribution capturing the mean and standard deviation of the observed VT durations. The prior represents the observer’s learned statistical knowledge of the VT duration distribution, i.e., the temporal context. From the posterior *p(VT*|*VT*_*m*_), the observer computes the Bayes least-squares (BLS) estimate as the posterior mean:

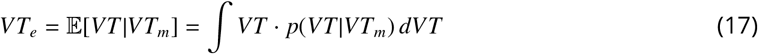

This minimizes the expected squared error 𝔼 *[(VT*_*e*_ − *VT)*^2^|*VT*_*m*_].

*Stage 3 (Response generation):* The estimated VT duration *VT*_*e*_ is transformed to predict the TTC. Since the occluded distance is half the visible distance (5 dva / 10 dva; Fig. 1A), a gain factor *G*_*f*_ = 0.5 is applied:

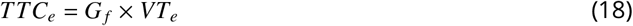

Finally, motor production noise is added to account for response variability: the actual TTC response is drawn from a scalar Gaussian distribution centered at (TTC_*e*_ with standard deviation *σ*_*p*_ = *w*_*p*_ × *TTC*_*e*_, where *w*_*p*_ is the Weber fraction for TTC production noise (scalar variability in motor output). To account for idiosyncratic response biases (e.g. consistently producing TTC responses earlier or later than expected), the model includes an offset parameter *o*_*b*_ (Remington et al., 2018).

Model parameters (*w*_*m*_, *w*_*p*_, *o*_*b*_) were fit by maximizing the log-likelihood of participants’ responses given the sampled VT durations using the *fminsearch* MATLAB (Mathworks) function. Simpson’s rule was used to numerically integrate the posterior during model fitting and simulation. To avoid local optima, parameter searches were repeated 20 times using different initialization values, showing a high degree of consistency between iterations. These model parameters were further used for model simulations, where we generated the same number of trials and TTC_*t*_ combinations as participants had in their dataset.

To evaluate our assumption that participants use both sources of information (Likelihood and Prior) to infer the TTC, we compared three model variants per participant (Fig. 4B): i) a Bayes Least-Squares (BLS) model that integrates both sources of information, with *w*_*m*_, *w*_*p*_ *and o*_*b*_ as free parameters; ii) a Maximum Likelihood Estimate (MLE) that ignores Prior information, with *w*_*m*_ as a free parameter, *w*_*p*_ fixed at the value obtained during the BLS fit, and *o*_*b*_ fixed at 0; iii) a Prior-dominant (PRI) model that ignores measurement-Likelihood information, with *w*_*p*_ fixed at the value obtained during the BLS fit, and *o*_*b*_ fixed at 0. Each model was fit separately to individual participants’ data, and model evidence was compared using the Bayesian Information Criterion (BIC) as a statistical metric. The BIC penalizes the use of excessive parameters: lower relative values indicate more parsimonious models that balance goodness-of-fit and simplicity to avoid overfitting. We expected the BLS model to show the lowest BIC within participants despite having more free parameters. To test this prediction, we calculated the ΔBIC between each alternative model and the BLS model and conducted one-tailed one-sample Wilcoxon signed-rank tests against 0 (Table S6A). Positive ΔBIC values indicate evidence in favor of the BLS model, while negative ΔBIC values indicate evidence in favor of the alternative model. The BLS model was statistically superior to the PRI (*µ*_Δ*BIC*_ = 2405.7) and the MLE models (*µ*_Δ*BIC*_ = 234.8).

Participants could improve TTC estimation either by learning the temporal context (as we assumed), or by increasing attention to the sampled durations. To disentangle which of these strategies could better explain participants’ behavior, we fit two variants of the BLS model separately to each run, and then evaluated changes in their parameters. Both model variants share the same underlying BLS architecture and total number of parameters; they differ only in which specific parameter was treated as free to account for behavioral changes across runs. We used as baseline parameters for each participant the values obtained from the previously described global fit of the BLS model to the full dataset (all four runs combined). In the Prior-learning model, *σ*_*prior*_ was the sole free parameter, while *w*_*m*_, *w*_*p*_, and *o*_*b*_ were constrained to the values obtained from the global fit. Conversely, in the Likelihood-sharpening model, only the measurement noise parameter *w*_*m*_ was free to vary, while *σ*_*prior*_ was fixed to the standard deviation of the observed VT durations and *w*_*p*_ and *o*_*b*_ were constrained to the global fit values. This approach allowed us to asses whether improvements in timing performance are better explained by the refinement of the prior (Fig. S5A, Upper left) or by reductions in measurement noise (Fig. S5A, Upper right), while keeping model complexity constant.

Performance improvements as a result of learning of the temporal context should be captured in the Prior-learning model by a decrease of the *σ*_*prior*_ parameter value across runs, reducing response variability at the expense of increased bias towards the mean (Fig. S5A, Bottom left). On the contrary, performance improvements due to increased attention to the sampled durations should be captured in the Likelihood-sharpening model by a decrease of the *w*_*m*_ parameter value across runs, reducing both response variability and bias towards the mean (Fig. S5A, Bottom right).

To test for differences in the relevant model parameters (MP _*j*_ ∈ {*σ*_*prior*_, *w*_*m*_}) across runs, we first ran a mixed-effects model with task run as predictor and separate intercepts per participant:

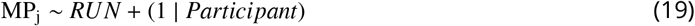

Considering that participants’ behavior reflected a bias towards the mean and in accordance with previous studies (Sohn, Lee, 2013), we expected a decrease in the *σ*_*prior*_ value across runs (Fig. S5B,C). *σ*_*prior*_ values differed across runs (Fig. S5B; MEM, *F*(3, 96) = 3.15, *p* = 0.028, *ϵ*^2^ = 0.06, *CI*: [0, 1]), and consistent with our prediction, post-hoc comparisons confirmed that *σ*_*prior*_ values decreased across runs in the Prior-learning model, ultimately converging towards the *σ* of the sampled durations (0.56 s; Fig. S5B; Table S7, one-tailed paired Wilcoxon signed-rank tests). In contrast, *w*_*m*_ values in the Likelihood-sharpening model remained constant across runs (Fig. S5C; MEM, *F*(3, 96) = 0.67, *p* = 0.572, *ϵ*^2^ = 0, *CI*: [0, 1]).

In line with the aforementioned behavioral bias, we further hypothesized that the Prior-learning model would be statistically superior to the Likelihood-sharpening model (i.e. would provide a better fit to participants’ data). To test this, for each participant we first summed the BIC values of each model fit across runs, calculated the Δ*BIC* between the two models and conducted a one-tailed one-sample Wilcoxon signed-rank test against 0. Positive Δ*BIC* values indicate evidence in favor of the Prior-learning model whereas negative values indicate evidence in favor of the Likelihood-sharpening (Fig. S5D).

To asses the BLS model performance in detail, we computed separately for each model and participant the average Variability and Bias across 100 simulations and compared it to the same metrics computed from participants’ behavior (Fig. 4C). We then computed the Mean Absolute Error (MAE) between observed and simulated response Variability and Bias, and compared it between models. To test our hypothesis that the BLS model had the lowest relative MAE we conducted one-tailed paired Wilcoxon signed-rank tests (Table S6B, C). From these simulated metrics (Variability and Bias) we also calculated the RMSE (RMSE_*model*_) and tested if the model could replicate the quadratic pattern across *TTC*_*t*_ observed in participants’ data (Fig. 1C, 4D). To do so we ran a mixed-effects model with RMSE_*model*_ as a dependent variable, TTC_*t*_ as a second-order orthogonal polynomial and separate intercepts and TTC_*t*_ slopes per participant:

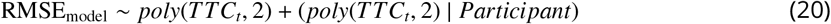

Furthermore, to test the model suitability to replicate the observed relationship between pmEC grid-like signals and RMSE (Fig. 3F), we used a mixed-effects model with RMSE_*model*_ as the dependent variable, and participants’ pmEC grid-like signal (GLS_*pmEC*_), target TTC (TTC_*t*_) and their interaction as predictors (4E). Separate intercepts and slopes were fit per participant:

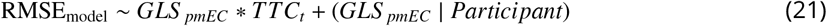

### Eye tracking

We used an MR-compatible infrared eye tracker with long-range optics (Eyelink 1000) to monitor gaze position at a rate of 500 hz during the experiment. After blink removal, the eye tracking data was linearly detrended, median centered, downsampled to the screen refresh rate of 120 hz and smoothed with a running-average kernel of 100 ms. Fixation error was computed separately for each participant and trial as the euclidean distance between the fixation target and the measured gaze position. In order to test for systematic imbalances or biases in viewing behavior, we partitioned participants’ fixation error for each target duration FE _*j*_ (where *j* ∈ {0.55, 0.67, 0.86, 1.2} s) an ran separate mixed-effect with FE _*j*_ as the dependent variable, target gaze direction as predictor, and participants as the error term:

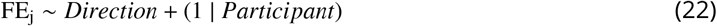

There were no significant differences in fixation error across directions (Fig. S2; Table S8).

## Supplementary Material

**Figure S1:**
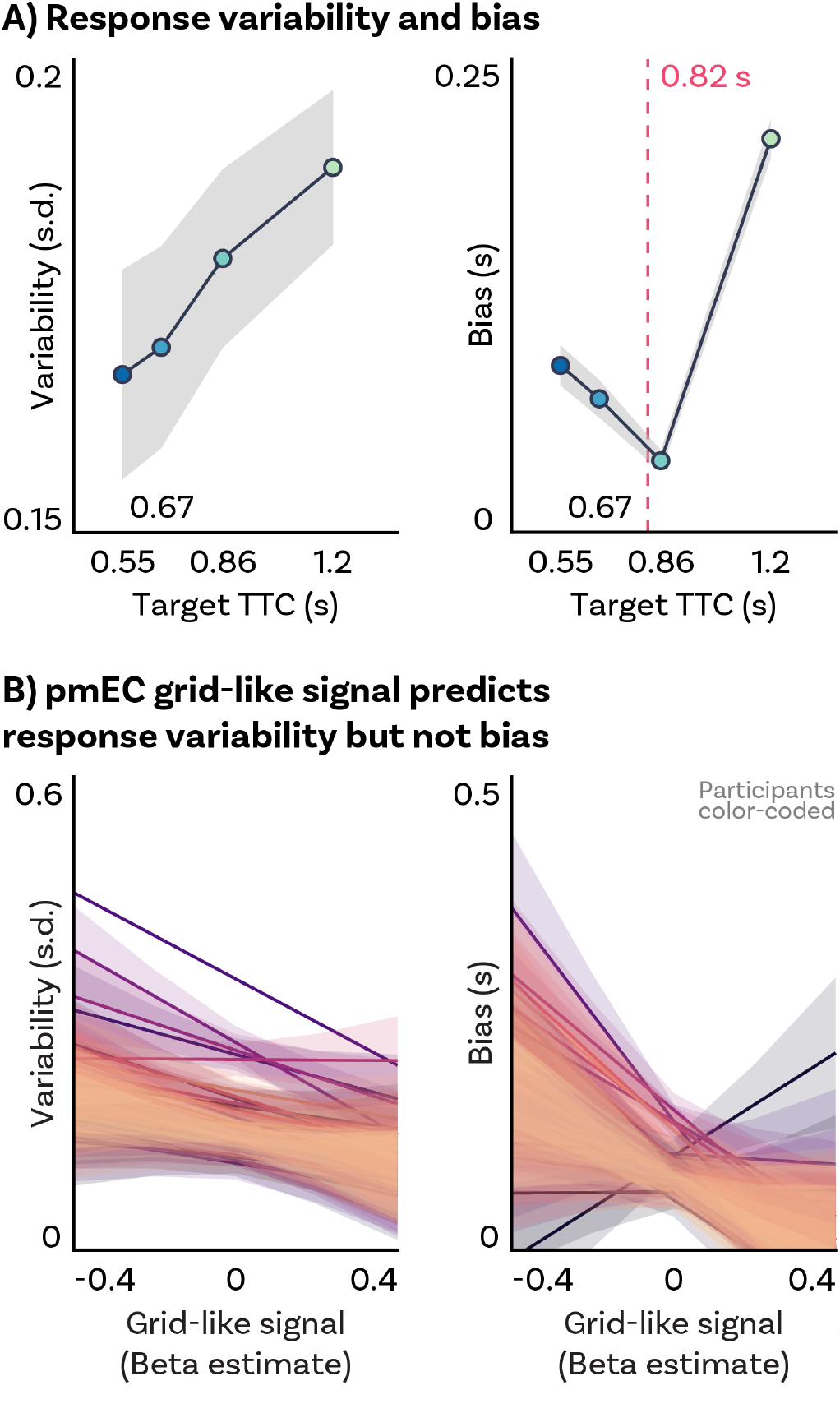
A) Response variability and bias. Left: Average response variability (standard deviation) across participants for each target TTC (TTC_*t*_). Response variability increases with longer TTC_*t*_. Right: Average response bias (error magnitude) across participants for each TTC_*t*_. Response bias follows a quadratic pattern across TTC_*t*_, increasing for durations further away from the mean of the sampled TTCs (magenta vertical dashed line). TTC_*t*_ are color coded. Group-level mean (colored circles) and SEM (gray shade). B) Within-subject pmEC grid-like modulation predicts response variability but not bias. After controlling for the effect of TTC_*t*_, stronger pmEC grid-like modulation is associated with lower response variability. Separate mixed-effect model (MEM) regression line fits and standard error shading are plotted for each participant (see Methods section for detailed description of the MEMs). Participants are color-coded

**Figure S2:**
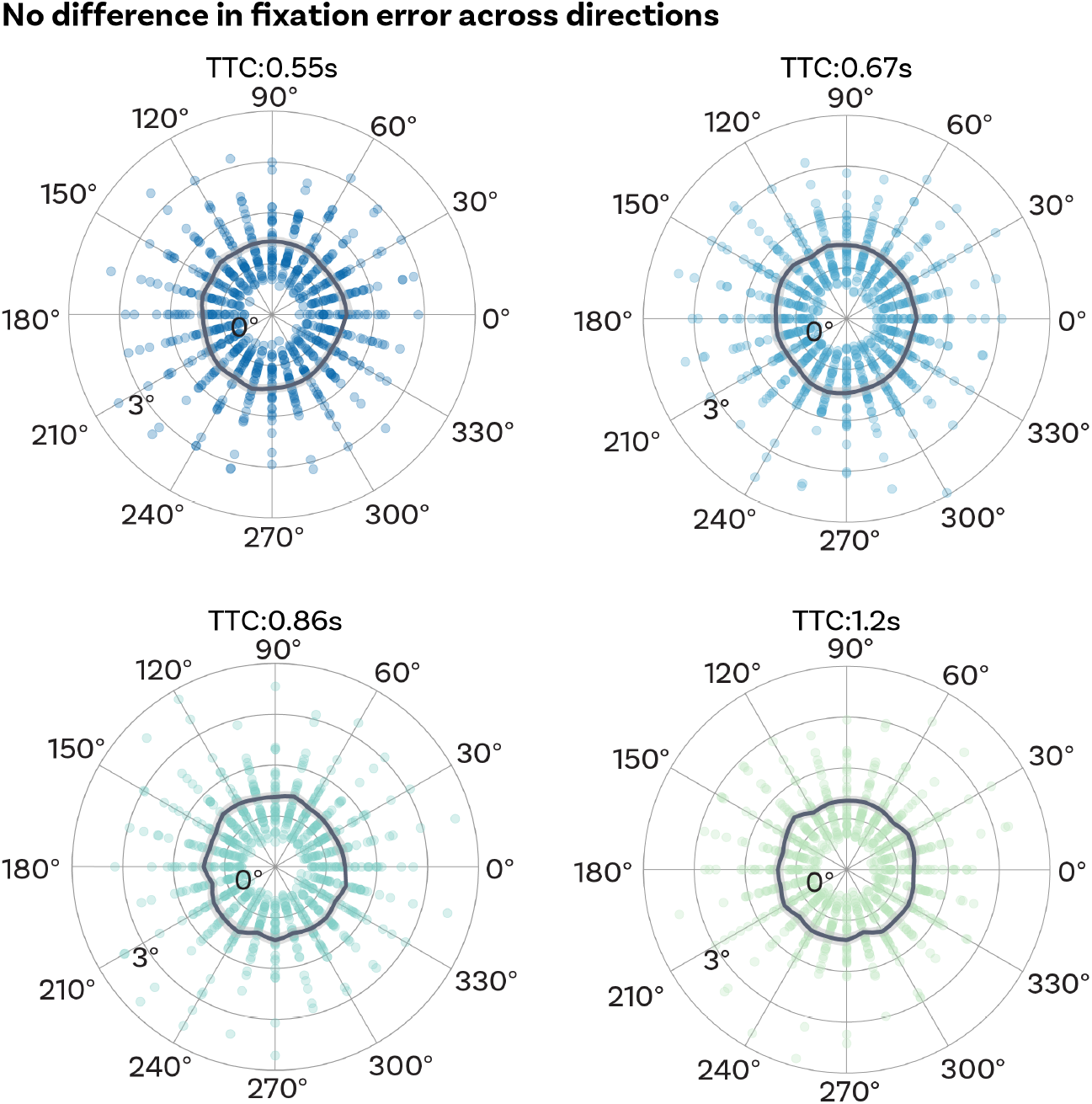
Euclidean distance between fixation target and gaze (fixation error). There were no significant differences in fixation error across all 24 gaze directions. Fixation quality does not affect the gaze-dependent hexadirectional modulation in EC presented in this study. Each dot per direction represents a single participant. Target TTCs are color coded. Group-level mean (black line) and SEM (gray shade). Fixation error displayed as degree of visual angle (radial axis). Single-participant data plotted as dots.

**Figure S3:**
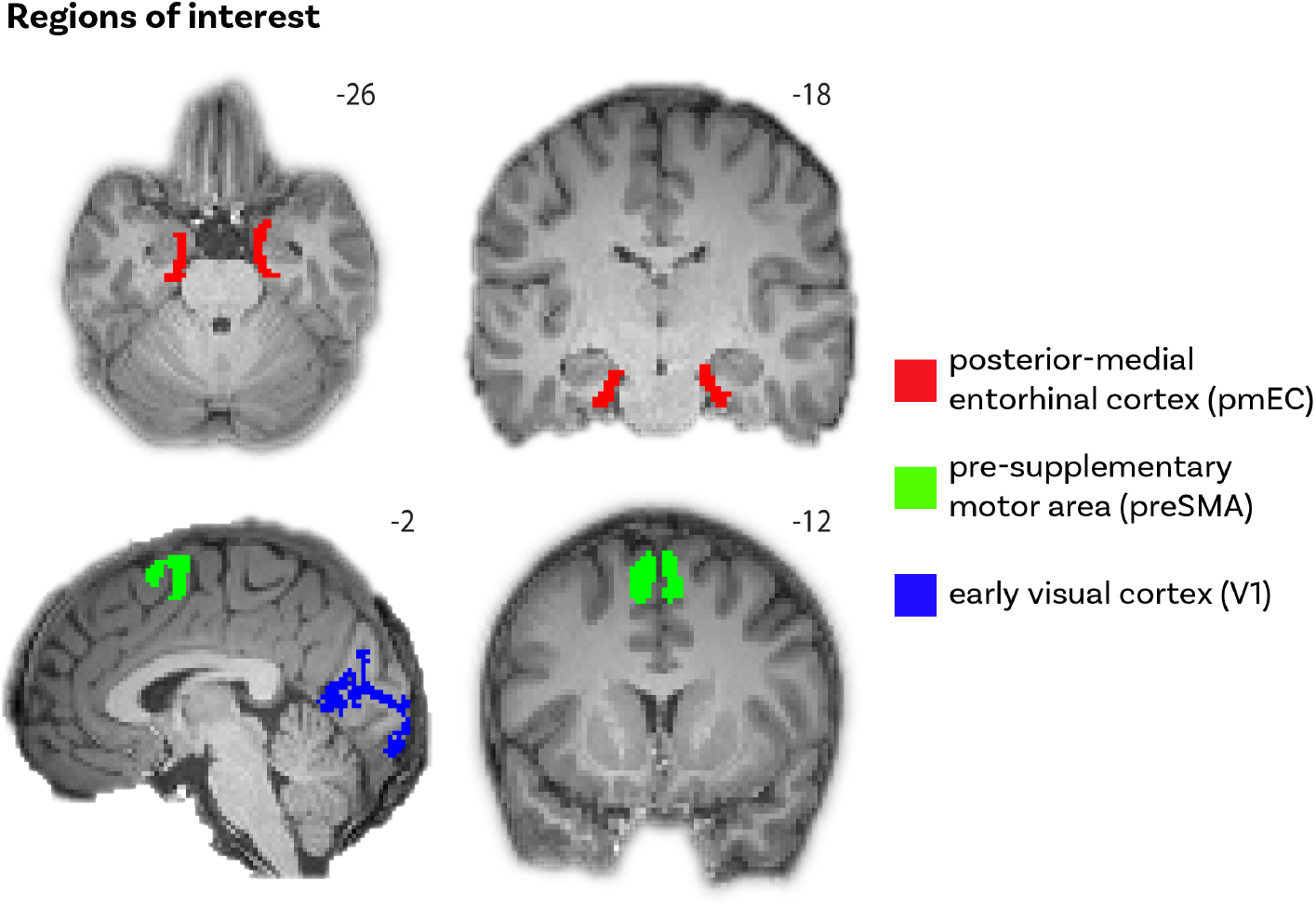
Regions of interest (ROIs). Red) posterior medial entorhinal cortex (pmEC) ROI representing the human homolog of rodent medial entorhinal cortex. This mask was obtained from Navarro Schröder et al. (2015). Green) pre-supplementary motor area (preSMA) obtained from the JuBrain SPM anatomy toolbox. Blue) early visual cortex (V1) anatomically defined for each participant using FreeSurfer’s cortical parcellation. ROIs superimposed onto a 2mm resolution skull-stripped structural template brain. MNI coordinates added.

**Figure S4:**
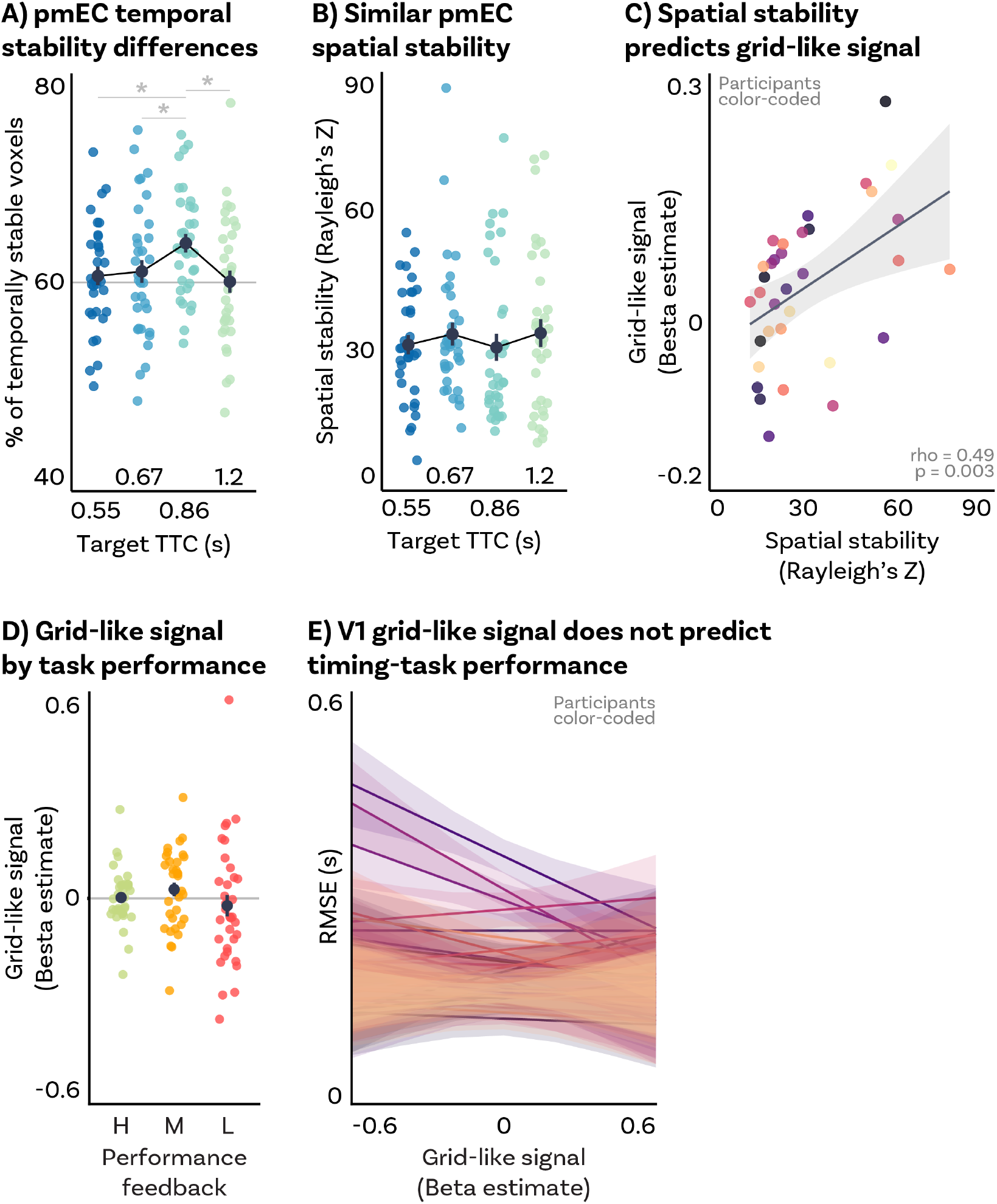
Additional pmEC grid-like signal analyses. A) Significant differences in temporal stability of grid-like signals in pmEC voxels across target TTCs. *TTC*_0.86_ showed the highest percent of pmEC temporally stable voxels. Target TTCs are color-coded. B) No significant difference in spatial stability of grid-like signals in pmEC voxels across target TTCs. Target TTCs are color-coded. C) *TTC*_0.86_ pmEC spatial stability predicts corresponding grid-like signal across participants. Each dot represents a single participant. Regression line (black) and standard error (gray shade). D) No significant pmEC grid-like signal modulation when grouping trials by task performance levels. No differences in pmEC grid-like signal modulation across task performance levels. Performance feedback levels are color-coded. E) Within-participant V1 grid-like signal does not predict TTC estimation RMSE. Separate regression lines are plotted for each participant. ABD) Depicted are the mean and SEM across participants (black dot and line) overlaid on single participant data (colored dots). CE) Participants are color-coded.

**Figure S5:**
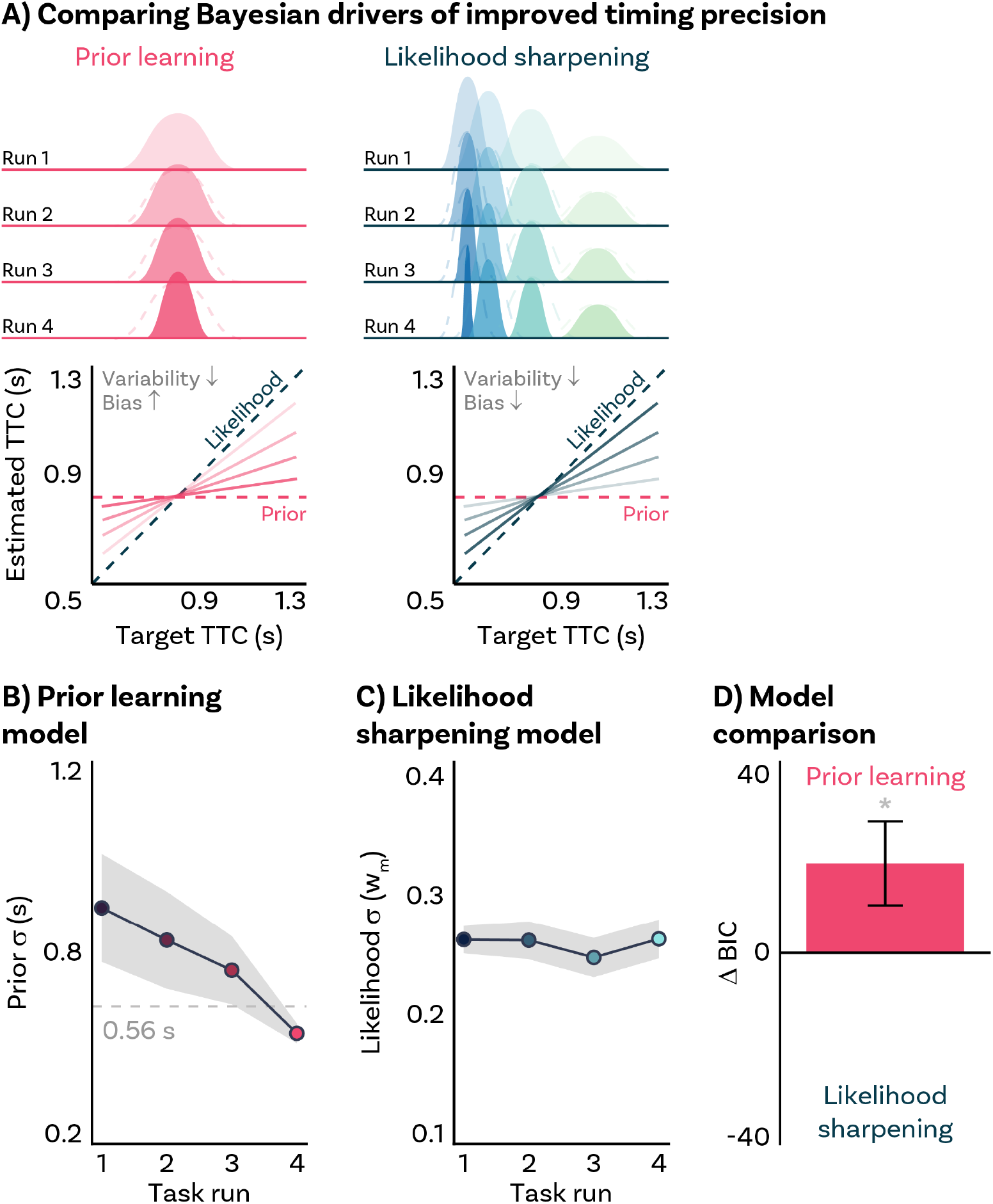
Comparison of Bayesian drivers of improved timing precision. A) Participants could improve their timing performance as a result of learning the temporal context (resulting in a Prior that decreases its standard deviation across runs; top left), or as a result of paying more attention to the sampled duration, thus reducing measurement noise (resulting in a Likelihood that decreases its standard deviation across runs; top right). A BLS observer with a more informative Prior will show a decrease in response variability at the expense of increased bias towards the mean (bottom left), whereas a BLS observer with a more informative Likelihood will show a decrease in both, response variability and bias (bottom right). Progression across runs is depicted by increasing color opacity. B) Prior-learning model. The *prior*_*σ*_ parameter value decreases across runs. C) Likelihood-sharpening model. The *w*_*m*_ parameter value remains stable across runs. D) Learning model comparison. Comparison of BIC values between models. A positive ΔBIC represents evidence in favor of the Prior-learning model as a better fit to participants’ data. BC) Group-level mean (colored circles) and SEM (gray shade). Runs are color-coded. D) Group-level mean (bar height) and SEM (black line). Statistics reflect p<0.05 (*) obtained using a group-level one-tailed one-sample Wilcoxon signed-rank test against zero.

**Table S1:** Independent ROI analysis for 6-fold symmetry in pmEC.

| One-tailed one-sample Wilcoxon signed-rank tests |  |  |  |  |  |  |
| --- | --- | --- | --- | --- | --- | --- |
| Target TTC | n | V statistic | p | $p_{FDR}$ | Effect size (r) | CI |
| 0.55 | 34 | 286 | 0.580 | 0.908 | -0.04 | [-0.35, 1] |
| 0.67 | 34 | 220 | 0.908 | 0.908 | -0.26 | [-0.53, 1] |
| 0.86 | 34 | 448 | 0.004 | 0.018 | 0.51 | [0.23, 1] |
| 1.2 | 34 | 224 | 0.896 | 0.908 | -0.25 | [-0.52, 1] |
| Two-tailed paired Wilcoxon signed-rank tests |  |  |  |  |  |  |
| Target TTC | n | V statistic | p | $p_{FDR}$ | Effect size (r) | CI |
| 0.86 vs. 0.55 | 34 | 425 | 0.029 | 0.023 | 0.43 | [0.07, 0.69] |
| 0.86 vs. 0.67 | 34 | 441 | 0.013 | 0.022 | 0.48 | [0.14, 0.72] |
| 0.86 vs. 1.2 | 34 | 439 | 0.014 | 0.022 | 0.48 | [0.13, 0.72] |

**Table S2:**
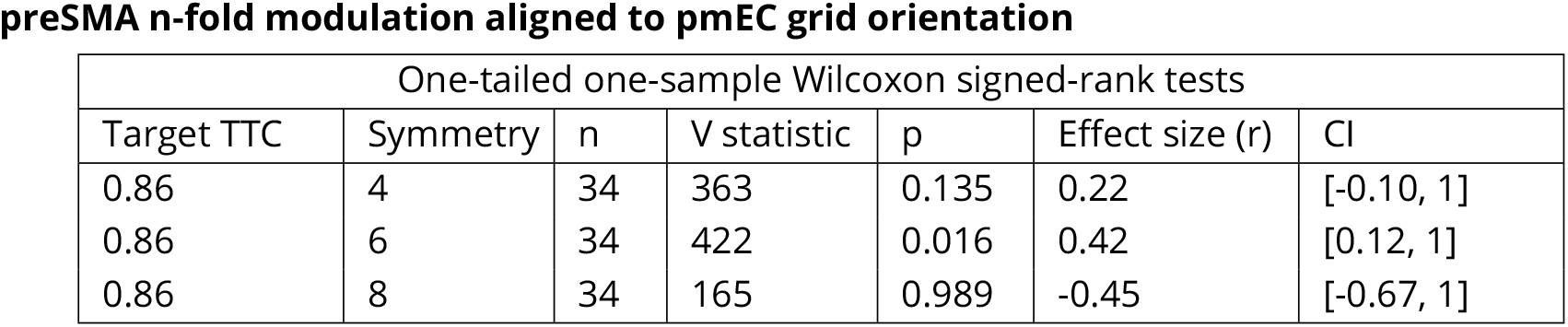
Independent ROI confirmatory analyses for 6-fold symmetry in preSMA.

**Table S3:**
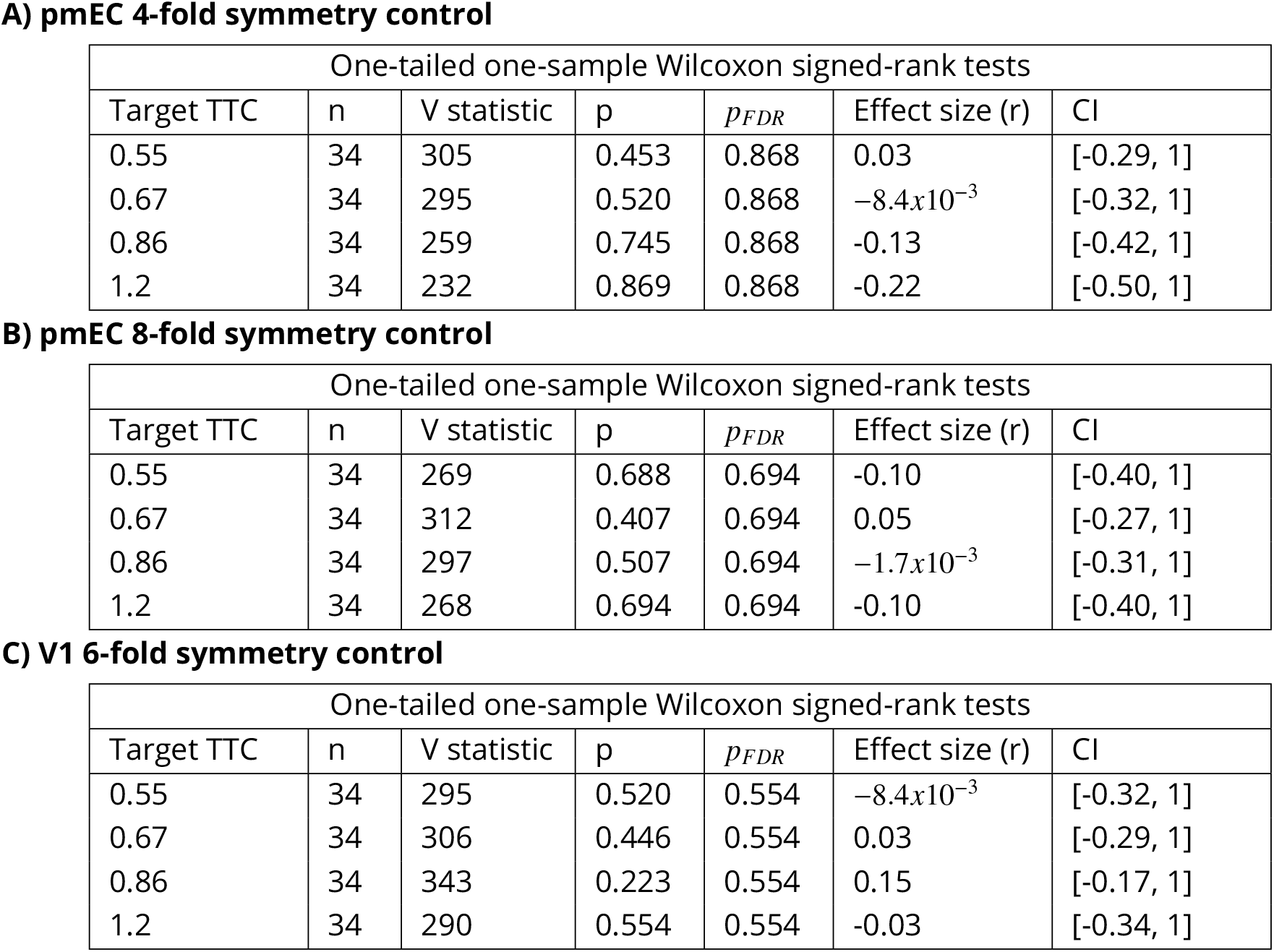
A) Independent ROI control analysis for 4-fold symmetry in pmEC. B) Independent ROI control analysis for 8-fold symmetry in pmEC. C) Independent ROI control analysis for 6-fold symmetry in V1.

**Table S4:** A) Clustering of grid orientations across pmEC voxels for each TTC_*t*_. B) Percent of pmEC temporally stable voxels for each TTC_*t*_.

**A) pmEC visual grid-like modulation spatial stability**
| One-tailed one-sample Wilcoxon signed-rank tests |  |  |  |  |  |  |
| --- | --- | --- | --- | --- | --- | --- |
| Target TTC | n | V statistic | p | $p_{FDR}$ | Effect size (r) | CI |
| 0.55 | 34 | 595 | $5.8 \times 10^{-11}$ | $5.8 \times 10^{-11}$ | 1 | [1, 1] |
| 0.67 | 34 | 595 | $5.8 \times 10^{-11}$ | $5.8 \times 10^{-11}$ | 1 | [1, 1] |
| 0.86 | 34 | 595 | $5.8 \times 10^{-11}$ | $5.8 \times 10^{-11}$ | 1 | [1, 1] |
| 1.2 | 34 | 595 | $5.8 \times 10^{-11}$ | $5.8 \times 10^{-11}$ | 1 | [1, 1] |

| One-tailed paired Wilcoxon signed-rank tests |  |  |  |  |  |  |
| --- | --- | --- | --- | --- | --- | --- |
| Target TTC | n | V statistic | p | $p_{FDR}$ | Effect size (r) | CI |
| 0.86 vs. 0.55 | 34 | 429 | 0.004 | 0.012 | 0.53 | [0.25, 1] |
| 0.86 vs. 0.67 | 34 | 337 | 0.042 | 0.042 | 0.36 | [0.05, 1] |
| 0.86 vs. 1.2 | 34 | 392 | 0.009 | 0.013 | 0.48 | [0.20, 1] |

**Table S5:** Independent ROI analysis for 6-fold symmetry in pmEC by behavioral performance level.

| One-tailed one-sample Wilcoxon signed-rank tests |  |  |  |  |  |  |
| --- | --- | --- | --- | --- | --- | --- |
| Performance level | n | V statistic | p | $p_{FDR}$ | Effect size (r) | CI |
| High | 34 | 299 | 0.493 | 0.739 | $5.04 \times 10^{-03}$ | [-0.31, 1] |
| Medium | 34 | 383 | 0.074 | 0.222 | 0.29 | [-0.03, 1] |
| Low | 34 | 233 | 0.865 | 0.865 | -0.22 | [-0.50, 1] |

**Table S6:**
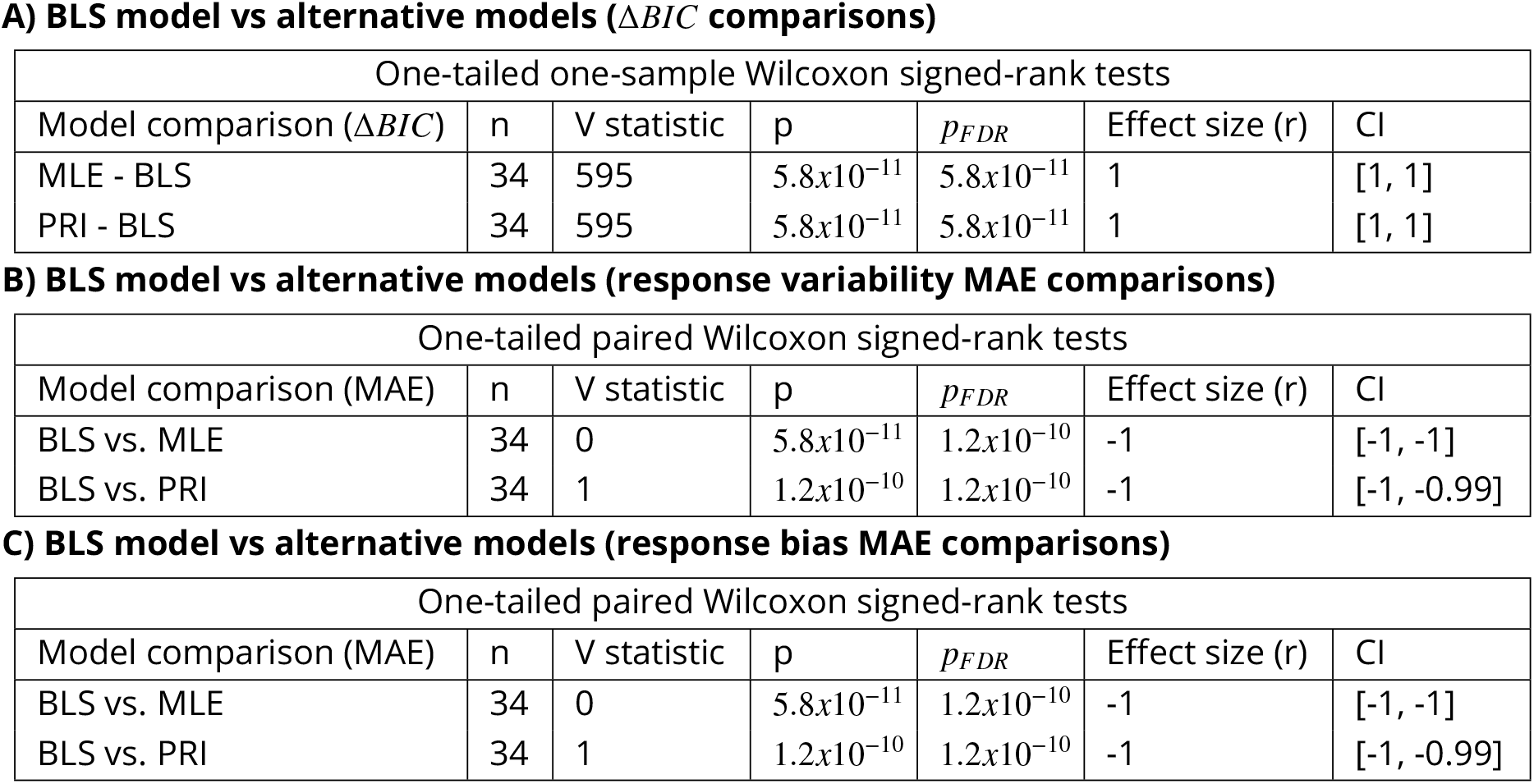
Comparison of BIC and MAE metrics between the BLS model and alternative models with different Likelihood / Prior informativeness.

**Table S7:**
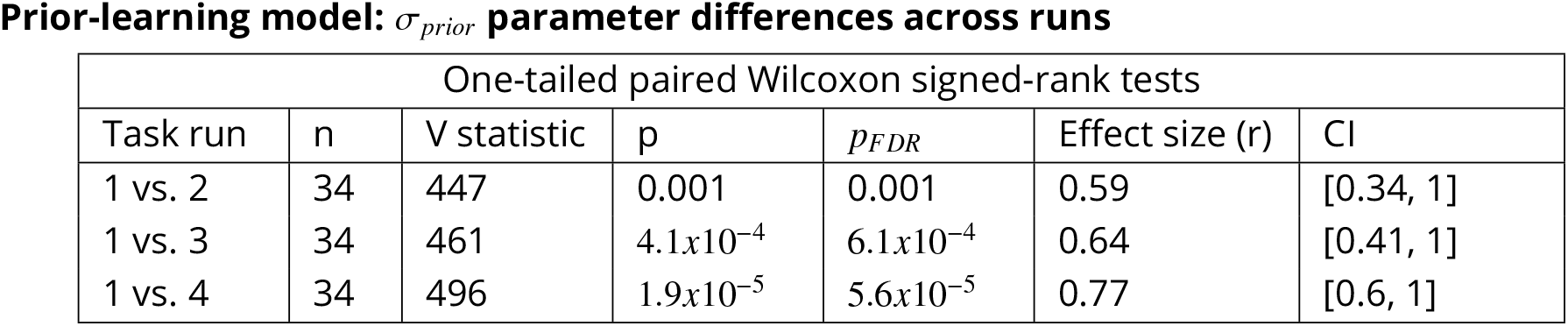
Prior-learning model. Post-hoc tests of *σ*_*prior*_ parameter value differences across runs.

**Table S8:** Fixation error across directions per target TTC.

| Mixed-Effect Model results |  |  |  |  |  |
| --- | --- | --- | --- | --- | --- |
| Target TTC (s) | DF | F | p-value | Effect size ($\epsilon^2$) | CI |
| 0.55 | 23 | 0.75 | 0.79 | $-7.5 \times 10^{-3}$ | [0.00, 0.00] |
| 0.67 | 23 | 1.33 | 0.14 | $9.6 \times 10^{-3}$ | [0.00, 0.00] |
| 0.86 | 23 | 1.31 | 0.15 | $9 \times 10^{-3}$ | [0.00, 0.00] |
| 1.2 | 23 | 0.98 | 0.48 | $-4.5 \times 10^{-4}$ | [0.00, 0.00] |

## References

Acerbi Luigi, Wolpert Daniel M., Vijayakumar Sethu. Internal Representations of Temporal Statistics and Feedback Calibrate Motor-Sensory Interval Timing // PLoS Computational Biology. 2012. 8, 11. e1002771.

Ambrogioni Luca, Ólafsdóttir H. Freyja. Rethinking the hippocampal cognitive map as a meta-learning computational module // Trends in Cognitive Sciences. 2023. 27, 8. 702–712.

Aronov Dmitriy, Nevers Rhino, Tank David W. Mapping of a non-spatial dimension by the hippocampal–entorhinal circuit // Nature. 2017. 543, 7647. 719–722.

Banino Andrea, Barry Caswell, Uria Benigno, Blundell Charles, Lillicrap Timothy, Mirowski Piotr, Pritzel Alexander, Chadwick Martin J., Degris Thomas, Modayil Joseph, Wayne Greg, Soyer Hubert, Viola Fabio, Zhang Brian, Goroshin Ross, Rabinowitz Neil, Pascanu Razvan, Beattie Charlie, Petersen Stig, Sadik Amir, Gaffney Stephen, King Helen, Kavukcuoglu Koray, Hassabis Demis, Hadsell Raia, Kumaran Dharshan. Vector-based navigation using grid-like representations in artificial agents // Nature. 2018. 557, 7705. 429–433.

Bao Xiaojun, Gjorgieva Eva, Shanahan Laura K., Howard James D., Kahnt Thorsten, Gottfried Jay A. Grid-like Neural Representations Support Olfactory Navigation of a Two-Dimensional Odor Space // Neuron. 2019. 102, 5. 1066–1075.e5.

Barry Caswell, Ginzberg Lin Lin, O’Keefe John, Burgess Neil. Grid cell firing patterns signal environmental novelty by expansion // Proceedings of the National Academy of Sciences of the United States of America. X 2012. 109, 43. 17687–17692.

Barry Caswell, Hayman Robin, Burgess Neil, Jeffery Kathryn J. Experience-dependent rescaling of entorhinal grids // Nature Neuroscience. VI 2007. 10, 6. 682–684. Number: 6 Publisher: Nature Publishing Group.

Behrens Timothy E.J., Muller Timothy H., Whittington James C.R., Mark Shirley, Baram Alon B., Stachenfeld Kimberly L., Kurth-Nelson Zeb. What Is a Cognitive Map? Organizing Knowledge for Flexible Behavior // Neuron. 2018. 100, 2. 490–509.

Bellmund Jacob, Gärdenfors Peter, Moser Edvard I., Doeller Christian F. Navigating cognition: Spatial codes for human thinking // Science. 2018. 362, 6415. eaat6766.

Bellmund Jacob, Polti Ignacio, Doeller Christian F. Sequence memory in the hippocampal–entorhinal region // Journal of Cognitive Neuroscience. 2020a. 32, 11. 2056–2070.

Bellmund Jacob L.S., Cothi William de, Ruiter Tom A., Nau Matthias, Barry Caswell, Doeller Christian F. Deforming the metric of cognitive maps distorts memory // Nature Human Behaviour. 2020b. 4, 2. 177–188.

Bellmund Jacob L.S., Deuker Lorena, Montijn Nicole D., Doeller Christian F. Mnemonic construction and representation of temporal structure in the hippocampal formation // Nature Communications. 2022. 13, 1. 3395.

Bellmund Jacob LS, Deuker Lorena, Doeller Christian F. Mapping sequence structure in the human lateral entorhinal cortex // eLife. VIII 2019. 8. e45333. Publisher: eLife Sciences Publications, Ltd.

Bicanski Andrej, Burgess Neil. Neuronal vector coding in spatial cognition // Nature Reviews Neuroscience. 2020. 21, 9. 453–470.

Bigus Erin R., Lee Hyun-Woo, Bowler John C., Shi Jiani, Heys James G. Medial entorhinal cortex mediates learning of context-dependent interval timing behavior // Nature Neuroscience. VIII 2024. 27, 8. 1587–1598. Publisher: Nature Publishing Group.

Boccara Charlotte N., Nardin Michele, Stella Federico, O’Neill Joseph, Csicsvari Jozsef. The entorhinal cognitive map is attracted to goals // Science. 2019. 363, 6434. 1443–1447.

Bousquet Olivier, Balakrishnan Karthik, Honavar Vasant. Is the Hippocampus a Kalman Filter? // Pacific Symposium on Biocomputing. 1998. 657–668.

Buckmaster Cindy A., Eichenbaum Howard, Amaral David G., Suzuki Wendy A., Rapp Peter R. Entorhinal Cortex Lesions Disrupt the Relational Organization of Memory in Monkeys // Journal of Neuroscience. XI 2004. 24, 44. 9811–9825. Publisher: Society for Neuroscience Section: Behavioral/Systems/Cognitive.

Bush Daniel, Barry Caswell, Manson Daniel, Burgess Neil. Using Grid Cells for Navigation // Neuron. 2015. 87, 3. 507–520.

Butler William N., Hardcastle Kiah, Giocomo Lisa M. Remembered reward locations restructure entorhinal spatial maps // Science. 2019. 363, 6434. 1447–1452.

Chang Chia Jung, Jazayeri Mehrdad. Integration of speed and time for estimating time to contact // Proceedings of the National Academy of Sciences of the United States of America. 2018. 115, 12. E2879–E2887.

Chen Xiaoli, He Qiliang, Kelly Jonathan W., Fiete Ila R., McNamara Timothy P. Bias in Human Path Integration Is Predicted by Properties of Grid Cells // Current biology: CB. 2015. 25, 13. 1771–1776.

Cheng Ken, Shettleworth Sara J., Huttenlocher Janellen, Rieser John J. Bayesian integration of spatial information // Psychological Bulletin. 2007. 133, 4. 625–637.

Cicchini Guido Marco, Arrighi Roberto, Cecchetti Luca, Giusti Marco, Burr David C. Optimal Encoding of Interval Timing in Expert Percussionists // Journal of Neuroscience. 2012. 32, 3. 1056–1060.

Clark Andy. Whatever next? Predictive brains, situated agents, and the future of cognitive science // Behavioral and Brain Sciences. 2013. 36, 3. 181–204.

Constantinescu Alexandra O., O’Reilly Jill X., Behrens Timothy E.J. Organizing conceptual knowledge in humans with a gridlike code // Science. 2016. 352, 6292. 1464–1468.

Cothi William de, Nyberg Nils, Griesbauer Eva-Maria, Ghanamé Carole Zisch Fiona, Lefort Julie M., Fletcher Lydia, Newton Coco, Renaudineau Sophie, Bendor Daniel, Grieves Roddy, Duvelle Éléonore, Barry Caswell, Spiers Hugo J. Predictive maps in rats and humans for spatial navigation // Current Biology. 2022. 32, 17. 3676–3689.e5.

Dayan Peter, Daw Nathaniel D. Decision theory, reinforcement learning, and the brain // Cognitive, Affective, & Behavioral Neuroscience. 2008. 8, 4. 429–453.

De Kock Rose, Zhou Weiwei, Joiner Wilsaan M, Wiener Martin. Slowing the body slows down time perception // eLife. IV 2021. 10. e63607. Publisher: eLife Sciences Publications, Ltd.

Dias Marcelo, Ferreira Raquel, Remondes Miguel. Medial Entorhinal Cortex Excitatory Neurons Are Necessary for Accurate Timing // Journal of Neuroscience. XII 2021. 41, 48. 9932–9943.

Doeller Christian F., Barry Caswell, Burgess Neil. Evidence for grid cells in a human memory network // Nature. 2010. 463, 7281. 657–661.

Doeller Christian F., King John A., Burgess Neil. Parallel striatal and hippocampal systems for landmarks and boundaries in spatial memory // Proceedings of the National Academy of Sciences. 2008. 105, 15. 5915–5920.

Duncan Stephen, Kuruvilla Maneesh V., Thompson Benjamin, Bush Daniel, Ainge James A. Grid cell distortion is associated with increased distance estimation error in polarized environments // Current Biology. X 2025. 35, 19. 4810– 4819.e5.

Eichenbaum Howard, Dudchenko Paul, Wood Emma, Shapiro Matthew, Tanila Heikki. The Hippocampus, Memory, and Place Cells: Is It Spatial Memory or a Memory Space? // Neuron. 1999. 23, 2. 209–226.

Epstein Russell A., Patai Eva Zita, Julian Joshua B., Spiers Hugo J. The cognitive map in humans: spatial navigation and beyond // Nature Neuroscience. 2017. 20, 11. 1504–1513.

Fernández Guillén, Brewer James B., Zhao Zuo, Glover Gary H., Gabrieli John D.E. Level of sustained entorhinal activity at study correlates with subsequent cued-recall performance: A functional magnetic resonance imaging study with high acquisition rate // Hippocampus. 1999. 9, 1. 35–44.

Fiser József, Berkes Pietro, Orbán Gergő, Lengyel Máté. Statistically optimal perception and learning: from behavior to neural representations // Trends in Cognitive Sciences. 2010. 14, 3. 119–130.

Fortin Norbert J., Agster Kara L., Eichenbaum Howard B. Critical role of the hippocampus in memory for sequences of events // Nature Neuroscience. 2002. 5, 5. 458–462.

Friston Karl, Buzsáki Gyorgy. The Functional Anatomy of Time: What and When in the Brain // Trends in Cognitive Sciences. 2016. 20, 7. 500–511.

Fuhs Mark C., Touretzky David S. Context Learning in the Rodent Hippocampus // Neural Computation. 2007. 19, 12. 3173–3215.

Garvert Mona M, Dolan Raymond J, Behrens Timothy EJ. A map of abstract relational knowledge in the human hippocampal–entorhinal cortex // eLife. 2017. 6. e17086.

Geerts Jesse P., Chersi Fabian, Stachenfeld Kimberly L., Burgess Neil. A general model of hippocampal and dorsal striatal learning and decision making // Proceedings of the National Academy of Sciences. 2020. 117, 49. 31427–31437.

Gibbon John. Scalar expectancy theory and Weber’s law in animal timing // Psychological Review. 1977. 84, 3. 279–325.

Graichen Luise P., Linder Magdalena S., Keuter Lars, Jensen Ole, Doeller Christian F., Lamm Claus, Staudigl Tobias, Wagner Isabella C. Entorhinal grid-like codes for visual space during memory formation // Nature Communications. X 2025. 16, 1. 9247.

Hafting Torkel, Fyhn Marianne, Molden Sturla, Moser May-Britt, Moser Edvard I. Microstructure of a spatial map in the entorhinal cortex // Nature. 2005. 436, 7052. 801–806.

Hahn Michael, Wei Xue-Xin. A unifying theory explains seemingly contradictory biases in perceptual estimation // Nature Neuroscience. IV 2024. 27, 4. 793–804. Publisher: Nature Publishing Group.

Hardcastle Kiah, Ganguli Surya, Giocomo Lisa M. Environmental Boundaries as an Error Correction Mechanism for Grid Cells // Neuron. V 2015. 86, 3. 827–839.

Hargreaves Eric L., Mattfeld Aaron T., Stark Craig E.L., Suzuki Wendy A. Conserved fMRI and LFP Signals during New Associative Learning in the Human and Macaque Monkey Medial Temporal Lobe // Neuron. 2012. 74, 4. 743–752.

Hartley Tom, Trinkler Iris, Burgess Neil. Geometric determinants of human spatial memory // Cognition. 2004. 94, 1. 39–75.

Heald James B., Wolpert Daniel M., Lengyel Máté. The Computational and Neural Bases of Context-Dependent Learning // Annual Review of Neuroscience. 2023. 46, 1. 233–258.

Henke Josephine, Bunk David, Werder Dina von, Häusler Stefan, Flanagin Virginia L, Thurley Kay. Distributed coding of duration in rodent prefrontal cortex during time reproduction // eLife. XII 2021. 10. e71612. Publisher: eLife Sciences Publications, Ltd.

Heys James G., Dombeck Daniel A. Evidence for a subcircuit in medial entorhinal cortex representing elapsed time during immobility // Nature Neuroscience. 2018. 21, 11. 1574–1582.

Heys James G., Rangarajan Krsna V., Dombeck Daniel A. The Functional Micro-organization of Grid Cells Revealed by Cellular-Resolution Imaging // Neuron. 2014. 84, 5. 1079–1090.

Heys James G., Wu Zihan, Allegra Mascaro Anna Letizia, Dombeck Daniel A. Inactivation of the Medial Entorhinal Cortex Selectively Disrupts Learning of Interval Timing // Cell Reports. 2020. 32, 12. 108163.

Hsieh Liang-Tien, Gruber Matthias J., Jenkins Lucas J., Ranganath Charan. Hippocampal activity patterns carry information about objects in temporal context // Neuron. 2014. 81, 5. 1165–1178.

Jazayeri Mehrdad, Shadlen Michael N. Temporal context calibrates interval timing // Nature Neuroscience. 2010. 13, 8. 1020–1026.

Jazayeri Mehrdad, Shadlen Michael N. A Neural Mechanism for Sensing and Reproducing a Time Interval // Current Biology. X 2015. 25, 20. 2599–2609.

Julian Joshua B., Doeller Christian F. Remapping and realignment in the human hippocampal formation predict context-dependent spatial behavior // Nature Neuroscience. VI 2021. 24, 6. 863–872.

Julian Joshua B., Keinath Alexandra T., Frazzetta Giulia, Epstein Russell A. Human entorhinal cortex represents visual space using a boundary-anchored grid // Nature Neuroscience. II 2018. 21, 2. 191–194.

Kang Yul Hr, Wolpert Daniel M., Lengyel Máté. Spatial uncertainty and environmental geometry in navigation // bioRxiv: The Preprint Server for Biology. 2023. 2023.01.30.526278.

Kanter Benjamin R., Lykken Christine M., Polti Ignacio, Moser May-Britt, Moser Edvard I. Event structure sculpts neural population dynamics in the lateral entorhinal cortex // Science. VI 2025. 388, 6754. eadr0927.

Keinath Alexandra T, Epstein Russell A, Balasubramanian Vijay. Environmental deformations dynamically shift the grid cell spatial metric // eLife. 2018. 7. e38169. Publisher: eLife Sciences Publications, Ltd.

Keinath Alexandra T., Rechnitz Ohad, Balasubramanian Vijay, Epstein Russell A. Environmental deformations dynamically shift human spatial memory // Hippocampus. 2021. 31, 1. 89–101.

Kessler Fabian, Frankenstein Julia, Rothkopf Constantin A. A Dynamic Bayesian Actor Model explains Endpoint Variability in Homing Tasks. 2022.

Killian Nathaniel J., Jutras Michael J., Buffalo Elizabeth a. A map of visual space in the primate entorhinal cortex // Nature. 2012. 491, 7426. 761–764.

Krupic Julija, Bauza Marius, Burton Stephen, Barry Caswell, O’Keefe John. Grid cell symmetry is shaped by environmental geometry // Nature. II 2015. 518, 7538. 232–235. Number: 7538 Publisher: Nature Publishing Group.

Kunz Lukas, Maidenbaum Shachar, Chen Dong, Wang Liang, Jacobs Joshua, Axmacher Nikolai. Mesoscopic Neural Representations in Spatial Navigation // Trends in Cognitive Sciences. 2019. 23, 7. 615–630.

Kunz Lukas, Schröder Tobias Navarro, Lee Hweeling, Montag Christian, Lachmann Bernd, Sariyska Rayna, Reuter Martin, Stirnberg Rüdiger, Stöcker Tony, Messing-Floeter Paul Christian, Fell Juergen, Doeller Christian F., Axmacher Nikolai. Reduced grid-cell–like representations in adults at genetic risk for Alzheimer’s disease // Science. 2015. 350, 6259. 430–433. Publisher: American Association for the Advancement of Science.

Körding Konrad P., Wolpert Daniel M. Bayesian integration in sensorimotor learning // Nature. 2004. 427, 6971. 244– 247.

Lisman John, Redish A.d. Prediction, sequences and the hippocampus // Philosophical Transactions of the Royal Society B: Biological Sciences. 2009. 364, 1521. 1193–1201. Publisher: Royal Society.

Maass Anne, Berron David, Libby Laura A, Ranganath Charan, Düzel Emrah. Functional subregions of the human entorhinal cortex // eLife. 2015. 4. e06426. Publisher: eLife Sciences Publications, Ltd.

Manns Joseph R., Eichenbaum Howard. Evolution of declarative memory // Hippocampus. 2006. 16, 9. 795–808.

Meirhaeghe Nicolas, Sohn Hansem, Jazayeri Mehrdad. A precise and adaptive neural mechanism for predictive temporal processing in the frontal cortex // Neuron. 2021. 109, 18. 2995–3011.e5.

Miyazaki Makoto, Nozaki Daichi, Nakajima Yasoichi. Testing Bayesian Models of Human Coincidence Timing // Journal of Neurophysiology. 2005. 94, 1. 395–399. Publisher: American Physiological Society.

Momennejad I., Russek E. M., Cheong J. H., Botvinick M. M., Daw N. D., Gershman S. J. The successor representation in human reinforcement learning // Nature Human Behaviour. 2017. 1, 9. 680–692.

Montchal Maria E., Reagh Zachariah M., Yassa Michael A. Precise temporal memories are supported by the lateral entorhinal cortex in humans // Nature Neuroscience. 2019. 22, 2. 284–288.

Moser Edvard I., Roudi Yasser, Witter Menno P., Kentros Clifford, Bonhoeffer Tobias, Moser May-Britt. Grid cells and cortical representation // Nature Reviews Neuroscience. 2014. 15, 7.

Nau Matthias, Julian Joshua B., Doeller Christian F. How the Brain’s Navigation System Shapes Our Visual Experience // Trends in Cognitive Sciences. 2018a. 22, 9. 810–825.

Nau Matthias, Navarro Schröder Tobias, Bellmund Jacob L. S., Doeller Christian F. Hexadirectional coding of visual space in human entorhinal cortex // Nature Neuroscience. 2018b. 21, 2. 188–190.

Nau Matthias, Schmid Alexandra C., Kaplan Simon M., Baker Chris I., Kravitz Dwight J. Centering cognitive neuroscience on task demands and generalization // Nature Neuroscience. IX 2024. 27, 9. 1656–1667.

Navarro Schröder Tobias, Haak Koen V, Zaragoza Jimenez Nestor I, Beckmann Christian F, Doeller Christian F. Functional topography of the human entorhinal cortex // eLife. VI 2015. 4. e06738. Publisher: eLife Sciences Publications, Ltd.

Nitsch Alexander, Garvert Mona M., Bellmund Jacob L. S., Schuck Nicolas W., Doeller Christian F. Grid-like entorhinal representation of an abstract value space during prospective decision making. 2023.

Niv Yael. Reinforcement learning in the brain // Journal of Mathematical Psychology. Special Issue: Dynamic Decision Making. 2009. 53, 3. 139–154.

O’Keefe J., Nadel L. The Hippocampus as a Cognitive Map. Oxford, UK: Oxford University Press, 1978.

Park Seongmin A., Miller Douglas S., Boorman Erie D. Inferences on a multidimensional social hierarchy use a grid-like code // Nature Neuroscience. 2021. 24, 9. 1292–1301.

Penny Will D., Zeidman Peter, Burgess Neil. Forward and Backward Inference in Spatial Cognition // PLOS Computational Biology. 2013. 9, 12. e1003383.

Peters-Founshtein Gregory, Dafni-Merom Amnon, Monsa Rotem, Arzy Shahar. Evidence for grid-cell-like activity in the time domain // Neuropsychologia. VI 2024. 198. 108878.

Petzschner Frederike H., Glasauer Stefan. Iterative Bayesian Estimation as an Explanation for Range and Regression Effects: A Study on Human Path Integration // Journal of Neuroscience. 2011. 31, 47. 17220–17229.

Petzschner Frederike H., Glasauer Stefan, Stephan Klaas E. A Bayesian perspective on magnitude estimation // Trends in Cognitive Sciences. 2015. 19, 5. 285–293.

Pezzulo Giovanni, Kemere Caleb, Meer Matthijs A.A. van der. Internally generated hippocampal sequences as a vantage point to probe future-oriented cognition // Annals of the New York Academy of Sciences. 2017. 1396, 1. 144–165.

Polti Ignacio, Nau Matthias, Kaplan Raphael, Wassenhove Virginie van, Doeller Christian F. Rapid encoding of task regularities in the human hippocampus guides sensorimotor timing // eLife. 2022. 11. e79027.

Remington Evan D., Parks Tiffany V., Jazayeri Mehrdad. Late Bayesian inference in mental transformations // Nature Communications. X 2018. 9, 1. 4419.

Rolando Felipe, Kononowicz Tadeusz W., Duhamel Jean-René, Doyère Valérie, Wirth Sylvia. Distinct neural adaptations to time demand in the striatum and the hippocampus // Current biology: CB. 2024. 34, 1. 156–170.e7.

Schapiro A., Turk-Browne N. Statistical Learning // Brain Mapping. 2015. 501–506.

Schapiro Anna C., Kustner Lauren V., Turk-Browne Nicholas B. Shaping of Object Representations in the Human Medial Temporal Lobe Based on Temporal Regularities // Current Biology. 2012. 22, 17. 1622–1627.

Schapiro Anna C., Turk-Browne Nicholas B., Norman Kenneth A., Botvinick Matthew M. Statistical learning of temporal community structure in the hippocampus // Hippocampus. 2016. 26, 1. 3–8.

Schiller Daniela, Eichenbaum Howard, Buffalo Elizabeth A., Davachi Lila, Foster David J., Leutgeb Stefan, Ranganath Charan. Memory and space: Towards an understanding of the cognitive map // Journal of Neuroscience. 2015. 35, 41. 13904–13911.

Sohn Hansem, Lee Sang-Hun. Dichotomy in perceptual learning of interval timing: calibration of mean accuracy and precision differ in specificity and time course // Journal of Neurophysiology. 2013. 109, 2. 344–362.

Sohn Hansem, Narain Devika, Meirhaeghe Nicolas, Jazayeri Mehrdad. Bayesian Computation through Cortical Latent Dynamics // Neuron. IX 2019. 103, 5. 934–947.e5.

Stachenfeld Kimberly L., Botvinick Matthew M., Gershman Samuel J. The hippocampus as a predictive map // Nature Neuroscience. XI 2017. 20, 11. 1643–1653.

Stangl Matthias, Achtzehn Johannes, Huber Karin, Dietrich Caroline, Tempelmann Claus, Wolbers Thomas. Compromised Grid-Cell-like Representations in Old Age as a Key Mechanism to Explain Age-Related Navigational Deficits // Current Biology. 2018. 28, 7. 1108–1115.e6.

Staresina Bernhard P., Alink Arjen, Kriegeskorte Nikolaus, Henson Richard N. Awake reactivation predicts memory in humans // Proceedings of the National Academy of Sciences. 2013. 110, 52. 21159–21164.

Staudigl Tobias, Leszczynski Marcin, Jacobs Joshua, Sheth Sameer A., Schroeder Charles E., Jensen Ole, Doeller Christian F. Hexadirectional Modulation of High-Frequency Electrophysiological Activity in the Human Anterior Medial Temporal Lobe Maps Visual Space // Current Biology. X 2018. 28, 20. 3325–3329.e4.

Syversen Ingrid Framås, Witter Menno P., Kobro-Flatmoen Asgeir, Goa Pål Erik, Navarro Schröder Tobias, Doeller Christian F. Structural connectivity-based segmentation of the human entorhinal cortex // NeuroImage. 2021. 245. 118723.

Theves Stephanie, Fernandez Guillén, Doeller Christian F. The Hippocampus Encodes Distances in Multidimensional Feature Space // Current Biology. 2019. 29, 7. 1226–1231.e3.

Theves Stephanie, Fernández Guillén, Doeller Christian F. The Hippocampus Maps Concept Space, Not Feature Space // Journal of Neuroscience. 2020. 40, 38. 7318–7325.

Tsao Albert, Sugar Jørgen, Lu Li, Wang Cheng, Knierim James J., Moser May-Britt, Moser Edvard I. Integrating time from experience in the lateral entorhinal cortex // Nature. IX 2018. 561, 7721. 57–62.

Vetter Philipp, Wolpert Daniel M. Context Estimation for Sensorimotor Control // Journal of Neurophysiology. 2000. 84, 2. 1026–1034.

Viganò Simone Bayramova Rena, Doeller Christian F., Bottini Roberto. Mental search of concepts is supported by egocentric vector representations and restructured grid maps // Nature Communications. 2023. 14, 1. 8132.

Viganò Simone Piazza Manuela. Distance and Direction Codes Underlie Navigation of a Novel Semantic Space in the Human Brain // Journal of Neuroscience. 2020. 40, 13. 2727–2736.

Viganò Simone Rubino Valerio, Soccio Antonio Di, Buiatti Marco, Piazza Manuela. Grid-like and distance codes for representing word meaning in the human brain // NeuroImage. 2021. 232. 117876.

Vo Annette, Tabrizi Nina S., Hunt Thomas, Cayanan Kayla, Chitale Saee, Anderson Lucy G., Tenney Sarah, White André O., Sabariego Marta, Hales Jena B. Medial entorhinal cortex lesions produce delay-dependent disruptions in memory for elapsed time // Neurobiology of Learning and Memory. 2021. 185. 107507.

Wagner Isabella C., Graichen Luise P., Todorova Boryana, Lüttig Andre, Omer David B., Stangl Matthias, Lamm Claus. Entorhinal grid-like codes and time-locked network dynamics track others navigating through space // Nature Communications. 2023. 14, 1. 231.

Wang Jing, Narain Devika, Hosseini Eghbal A., Jazayeri Mehrdad. Flexible timing by temporal scaling of cortical responses // Nature Neuroscience. I 2018. 21, 1. 102–110. Number: 1 Publisher: Nature Publishing Group.

Wang Jingyi, Tambini Arielle, Pritschet Laura, Taylor Caitlin M., Jacobs Emily G., Lapate Regina C. The intrinsic time tracker: temporal context is embedded in entorhinal and hippocampal functional connectivity patterns // Nature Communications. X 2025. 16, 1. 8817.

Whittington James C. R., Muller Timothy H., Mark Shirley, Chen Guifen, Barry Caswell, Burgess Neil, Behrens Timothy E. J. The Tolman-Eichenbaum Machine: Unifying Space and Relational Memory through Generalization in the Hippocampal Formation // Cell. XI 2020. 183, 5. 1249–1263.e23.

Wiener Martin, Michaelis Kelly, Thompson James C. Functional correlates of likelihood and prior representations in a virtual distance task // Human Brain Mapping. 2016. 37, 9. 3172–3187.

Witter Menno P., Amaral David G. CHAPTER 21 - Hippocampal Formation // The Rat Nervous System (Third Edition). Burlington: Academic Press, I 2004. 635–704.

World Medical Association. World Medical Association Declaration of Helsinki: Ethical Principles for Medical Research Involving Human Subjects // JAMA. 2013. 310, 20. 2191–2194.

